# Central dogma rates and the trade-off between precision and economy

**DOI:** 10.1101/276139

**Authors:** Jean Hausser, Avi Mayo, Uri Alon

## Abstract

Steady-state protein abundance is set by four rates: transcription, translation, mRNA decay and protein decay. A given protein abundance can be obtained from infinitely many combinations of these rates. This raises the question of whether the natural rates for each gene result from historical accidents, or are there rules that give certain combinations a selective advantage? We address this question using high-throughput measurements in rapidly growing cells from diverse organisms to find that about half of the rate combinations do not exist: genes that combine high transcription with low translation are strongly depleted. This depletion is due to a trade-off between precision and economy: high transcription decreases stochastic fluctuations but increases transcription costs. Our theory quantitatively explains which rate combinations are missing, and predicts the curvature of the fitness function for each gene. It may guide the design of gene circuits with desired expression levels and noise.

## Introduction

To function well in a given environment, cells need to express genes at the right protein copy number^14, 41,67^. Steady-state protein abundance is set by two reactions of synthesis —transcription and translation —balanced by two processes of removal – dilution / degradation of mRNAs and proteins^27^. Together, these make up the four basic rates of the central dogma^12^.

The rates of these four central dogma reactions are controlled by diverse regulators. Transcription rate is set by transcription factors and chromatin remodelers ^40^. Translation is modulated by RNA binding proteins and non-coding RNAs^36,65^, and so on. The effects of these molecular controls can be summarized by the central dogma rates, such that each protein is a point in a four dimensional space whose axes are the four rates. In this study, we name this the ‘Crick space’, in honor of Francis Crick who proposed the central dogma^12^.

One important property of Crick space is that the same steady-state protein abundance can be achieved by many combinations of rates. For example, consider a protein made at 1000 copies per hour (Fig. 1). This can be achieved by transcribing 100 mRNAs and translating 10 proteins from each mRNA every hour. Alternatively, the 1000 proteins could be made from one mRNA translated into 1000 proteins per hour (in this example, we fixed mRNA and protein decay). There is an infinite number of ways to combine transcription and translation rates, *β*_*m*_ and *β*_*p*_, in order to supply a given steady-state number of proteins *p*, namely *β*_*m*_*β*_p_/*α*_m_*α*_*p*_ = *p* where *α*_*m*_ and *α*_*p*_ are the mRNA and protein decay rates.

**Figure 1.**
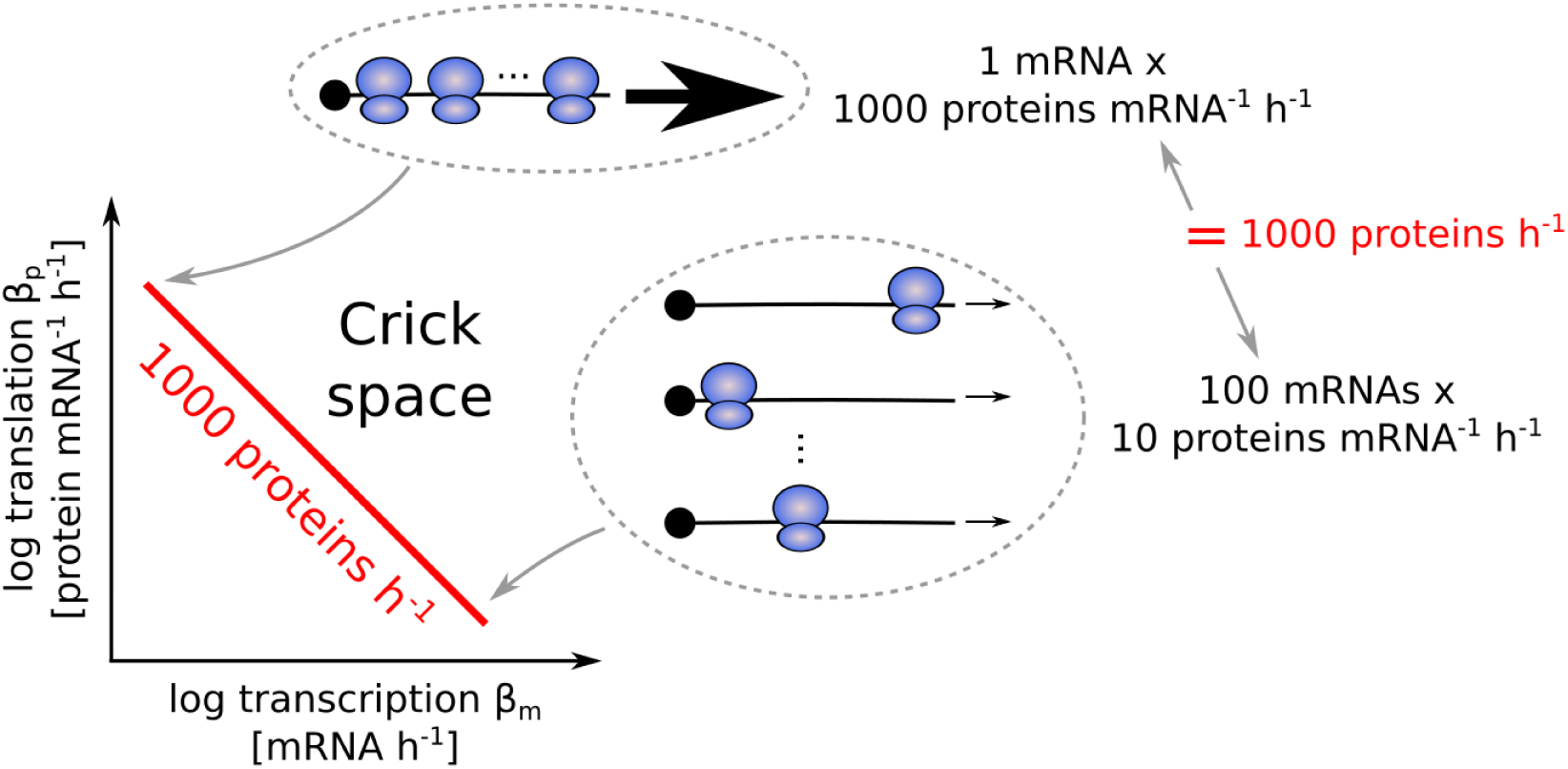
For each gene, an infinite number of combinations of transcription and translation rates can achieve a given protein abundance. For example, to obtain 1000 proteins per hour, one possibility is to translate 1000 proteins per hour from a single mRNA. Another option is to translate 10 proteins per hour from 100 mRNAs. We assumed fixed mRNA and protein decay to simplify the visualization. See also Figure S1.

Here we ask whether such combinations occur randomly, as expected if they are equally beneficial and historical accident or genetic drift is at play, or whether there are rules based on specific translation / transcription ratios that have selective advantage. If such rules exist, we might expect to see patterns in the way that genes occupy Crick space.

While there has not been systematic evidence for rules so far, previous work described how different combinations of central dogma rates can differ in their biological impact. One line of work shows that intrinsic noise^8,17,57,60^, the stochastic variation in protein number due to small-number effects, is largest when there are few mRNAs translated into many proteins^5,45,52,54^. This large noise occurs because the relative fluctuations in the number of a few mRNAs are large, and are amplified by strong translation. This idea was first proposed by McAdams and Arkin ^45^ based on theoretical arguments. The prediction that a given protein abundance can be reached with the least noise when transcription is high and translation is low was validated by Ozbudak et al. ^52^ using synthetic constructs with defined transcription and translation rates. Bar-Even et al. ^5^ measured the noise and abundance of 43 *S. cerevisiae* proteins and found that noise scaled with protein abundance in a way consistent with the predictions of McAdams and Arkin ^45^. Also in *S. cerevisiae*, Newman et al. ^50^ observed a correlation between noise and mRNA abundance for 2500+ genes.

Another difference between combinations of rates that give the same steady-state protein abundance is mRNA cost — the reduction in fitness due to production of mRNA^24,31,44,59,82^. Theoretical studies proposed that the cost of synthesizing mRNAs confers a selectable disadvantage^45,59,82^. In *S. cerevisiae*, expressing a non-beneficial mRNA penalizes the growth rate in proportion to the transcription rate^31^. In *E. coli*, expressing a protein at a given abundance from a larger number of mRNAs decreases fitness^24^.

Noise and cost are thought to be significant components in determining the fitness and selection of biological designs^20,32,39,45,66,81^. In particular, McAdams and Arkin ^45^ and others^52^ proposed that there should be a trade-off between minimizing gene expression costs and minimizing noise in protein abundance. Testing this hypothesis has been difficult, partly because the central dogma rates could not be measured genome-wide until recently^43^. Another hurdle that prevented testing the hypothesis of a precision - economy trade-off in gene expression is that it is unclear how the interplay of precision and economy should affect the distribution of genes in the Crick space.

Here we address the question of rules for protein expression by analyzing comprehensive data on central dogma rates from several model organisms^16,41,49,50,66,84^ and by theory on evolutionary trade-offs. We find that about half of the Crick space is empty: genes do not seem to combine high transcription with low translation. This depleted region is accessible by synthetic constructs, and hence its emptiness is not based on mechanistic constraints. We explain the empty Crick space by a trade-off between cost and noise of gene expression. This theory accurately predicts the boundary of the empty region which varies by 2 orders of magnitude between the model organisms we considered. This approach might be of use to design synthetic gene expression circuits, and suggests rules for central dogma rates that seem to apply from bacteria to humans.

## Results

### Genes combining high transcription and low translation are depleted

We estimated transcription *β*_*m*_ and translation *β*_*p*_ rates, as well as mRNA and protein decay rates, for thousands of genes from previous mRNAseq and ribosome profiling (RP) experiments in *S. cerevisiae*^84^, *M. musculus, H. sapiens*^16^ and *E. coli*^41^(Methods). All data were collected under conditions of rapid growth.

We find, in accordance with previous studies^30,41,42,66^, that transcription and translation rates in rapidly0020growing cells vary much more from gene to gene than mRNA and protein decay rates. Transcription and translation rates vary over a 1000-fold range compared to a 10-fold range for decay rates of mRNA and protein (Fig. S1A–D). We therefore simplify our discussion by considering a 2D Crick space, formed by transcription and translation rates.

Reducing the 4-dimensional Crick space to two-dimensions neglects aspects of cell biology such as the dynamics of gene regulation in response to environmental perturbations^56,80^, but it allows us to focus on the most variable rates in setting steady-state protein abundance in growing cells, transcription and translation, and ask what rules may underlie them.

Plotting the transcription and translation rates of genes in four model organisms, we observe common boundaries in the Crick space (Fig. 2). First, the maximal translation is 10^3.6^ – 10^4^ proteins per mRNA per hour, a bound that can be explained from the ribosome translocation speed (Methods). Second, we observed lower bounds on the transcription rate and on the product of transcription and translation. These boundaries stem from technical limits of the assays (Fig. S2A).

**Figure 2.**
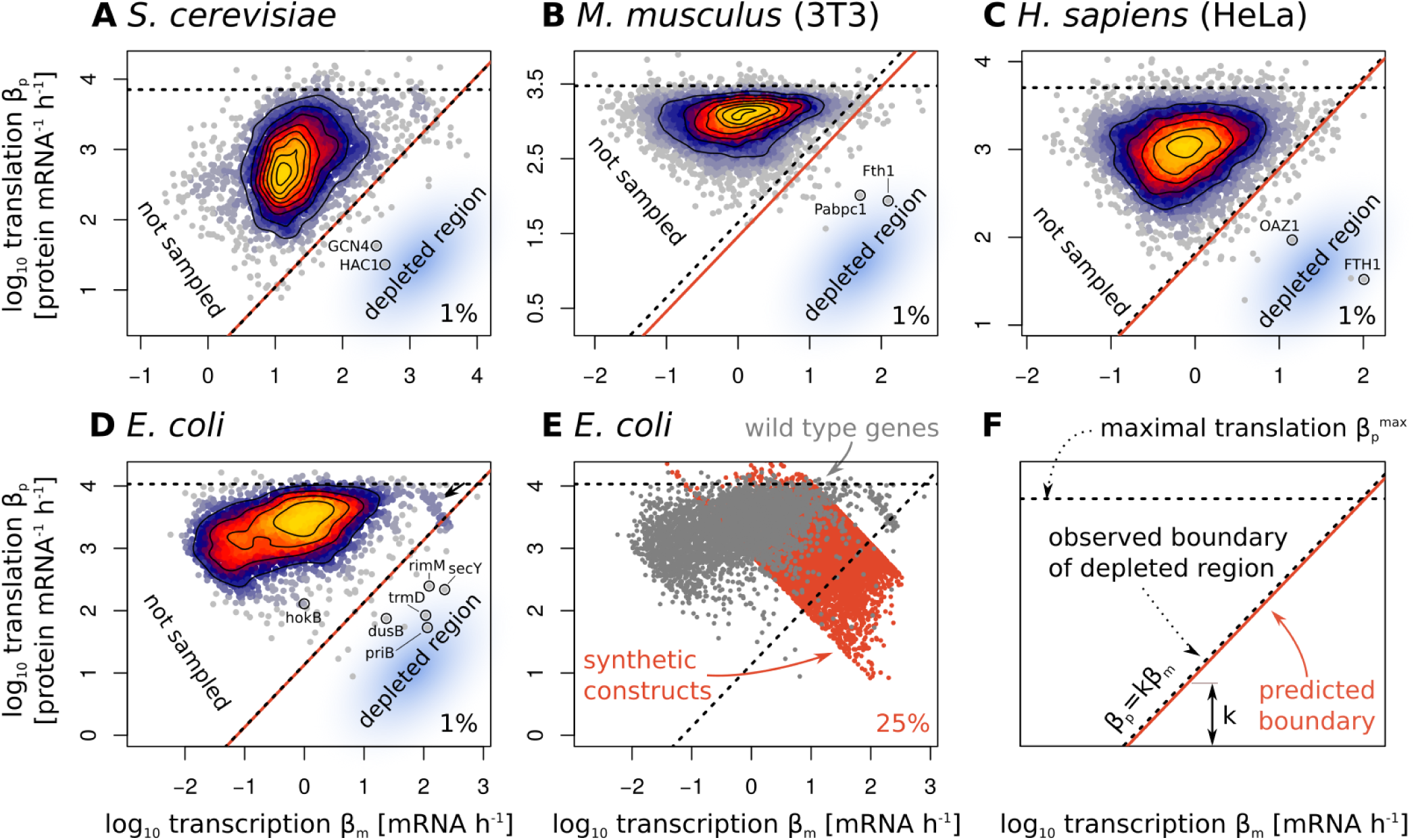
Genes combining high transcription and low translation are depleted from the Crick space across organisms. **A–D.** There is a depleted region in the Crick space of four model organisms. Transcription and translation rates were estimated from ribosome profiling and mRNA sequencing data. The top percentile of translation rates 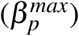 is represented as a horizontal dashed line. The observed boundary of the depleted region (diagonal dashed line) has slope 1 and is such that 99% of the genes have a larger translation / transcription ratio. Excluding 1% of genes in this way makes the boundary line less sensitive to measurement errors and to outlier genes, some of which are highlighted. The predicted boundary of the depleted region (red line) is according to the theory introduced later in this article. Technical constraints explain the absence of genes at low transcription and translation rates (region marked ‘not sampled’). *S. cerevisiae* data (**A**) from Weinberg et al. ^84^. *M. musculus* (**B**) and *H. sapiens* (**C**) data from Eichhorn et al. ^16^. *E. coli* data (**D**) from Li et al. ^41^. **E.** Transcription and translation rates of 3744 *E. coli* genes (gray dots) and of 7624 synthetic constructs (red dots) of Kosuri et al. ^37^. The apparent negative correlation between transcription and translation rates in this dataset is due to limits in the linear range of flow cytometry measurements which leads to censoring of low and high abundance proteins^37^. **F.** Figure legend summarizing the meaning of the different lines on panels A–E. See also Figure S2.

Unexpectedly, and most importantly for the present study, there was a lack of genes combining high transcription with low translation (blue regions in Fig. 2). We call this region the depleted region of the Crick space. This depleted region makes up about half of the Crick space and is bounded by a line of constant ratio between transcription *β*_*m*_ and translation *β*_*p*_, *β*_*p*_/*β*_*m*_ = *k*, with *k* = 1.1 *±* 0.1, 14 *±* 3, 44 *±* 3, 66 *±* 4 in *S. cerevisiae, E. coli, M. musculus* and *H. sapiens*, respectively (Table 1). In logarithmic axes, the boundary of the depleted region has slope 1 and intercept log(*k*). Hence, *k* determines the boundary of the depleted region.

**Table 1.** The intercept *k* of the boundary of the depleted region varies over two orders of magnitude across the studied organisms. *k* can be predicted from the maximal translation rate 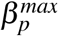 [protein mRNA^−1^ *h*^−1^], the protein decay rate *α*_*p*_ [h^−1^], the total transcriptional output Σ *β*_*m*_ [mRNA h^−1^], and the noise floor *c*_*v*0_ using Eq. 5. The measured *k* is defined by having 99% of genes with *β*_*p*_/*β*_*m*_ > *k*, with error bars from bootstrapping. Error bars indicate the standard error.

| Organism | measured $k$ | predicted $k$ | $\beta_p^{max}(\times 10^3)$ | $\alpha_p$ | $\Sigma \beta_m(\times 10^4)$ | $c_{v0}$ |
| --- | --- | --- | --- | --- | --- | --- |
| *S. cerevisiae* | $1.1 \pm 0.1$ | $1.1 \pm 0.3$ | $7.1 \pm 0.7$ | $1.3 \pm 0.2$ | $30 \pm 15$ | $0.10 \pm 0.01$ |
| *E. coli* | $14 \pm 3$ | $13 \pm 4$ | $11 \pm 1$ | $1.9 \pm 0.3$ | $2.0 \pm 1$ | $0.25 \pm 0.01$ |
| *M. musculus* (3T3) | $44 \pm 3$ | $29 \pm 8$ | $3.0 \pm 0.3$ | $0.04 \pm 0.01$ | $2.5 \pm 1.2$ | $0.3 \pm 0.03$ |
| *H. sapiens* (HeLa) | $66 \pm 4$ | $60 \pm 17$ | $5.1 \pm 0.5$ | $0.05 \pm 0.01$ | $1.4 \pm 0.7$ | $0.3 \pm 0.03$ |

In *E. coli*, the boundary of the depleted region has additional structure. Departing from the main distribution of genes are a set of 59 genes with very high transcription and translation rates (*β*_*m*_ *>* 80 h^−1^ and *β*_*p*_ *>* 1000 mRNA^−1^ h^−1^, indicated by an arrow in Fig. 2D). 90% of these genes are ribosomal proteins. The rest are high-abundance proteins such as the glycolysis enzyme *gapA* (the *E. coli* equivalent to *Gapdh*), the ATP synthase c subunit, and outer-membrane proteins (OMPs). Most genes in this set are essential to cellular viability, as indicated by knock-out experiments (Fig. S2B). If one removes essential genes from the data, the boundary of the depleted region shifts up and tightly fits the rest of the genes with a higher intercept, *k* = 44 *±* 9. This higher intercept for non-essential genes is predicted by the theory introduced below (Fig. S2B, Methods).

A depleted region is also found when we estimate transcription and translation from two proteomics and mRNAseq datasets in *H. sapiens* and *M. musculus* (Fig. S2C–D). Finally, we observe the depleted region when plotting the transcription burst rate against the translational burst size^22,23^ inferred from single cell protein abundance measurements^50^ (Fig. S2E).

We assessed the statistical significance of the depleted region by shuffling transcription and translation rates while conserving the distributions of protein abundance and translation rates (Fig. S2F–H). We find that none of the 10^4^ shuffled datasets show a comparable depleted region in Crick space (equal or smaller number of genes with *β*_*p*_/*β*_*m*_ *< k, p <* 10^−4^).

### Genes can mechanistically achieve high transcription and low translation

A possible explanation for the depleted region is that a (possibly yet unknown) biochemical constraint prevents high transcription combined with low translation. To test for this possibility, we re-analyzed measurements by Kosuri et al. ^37^ on synthetic genes that provided a wide range of transcription and translation rates. In that study, GFP was expressed in *E. coli* under the control of 114 promoters and 111 Ribosomal Binding Sites (RBSs) of varying strengths. Relative abundance of the GFP mRNA and protein was then quantified by mRNAseq and flow cytometry.

The transcription and translation rates of the synthetic constructs largely overlap with those of *E. coli* genes (Fig. 2E). However, in contrast to *E. coli* genes, a large fraction of the synthetic constructs achieved a combination of high transcription and low translation rates. 25% of the synthetic genes fall in the depleted region seen for endogenous *E. coli* genes.

This observation supports the conclusion that the biochemistry of gene expression can achieve high transcription and l006Fw translation in principle. In support of this argument, there are indeed examples of such genes in the depleted region for all four organisms. These include the ribosome maturation factor rimM in *E. coli*, the amino acid response regulator GCN4 in *S. cerevisiae*, and the iron homeostasis protein Fth1 in *M. musculus* and *H. sapiens* (Fig. 2A–D). Thus, the results in this paper concern *∼* 99% of the genes, with the remaining *∼* 1% requiring additional analysis (see discussion for suggested effects for these genes).

### Increasing transcription at constant protein abundance increases transcriptional cost while decreasing stochastic fluctuations

Because biochemical constraints do not seem to explain the lack of genes combining high transcription with low translation, we asked whether evolutionary trade-offs might explain it.

One could hypothesize that cells avoid combining high transcription with low translation in order to minimize the cost of mRNA synthesis. In *S. cerevisiae* growing in rich medium, the fitness cost of mRNA is *c*_*m*_ *∼* 10^−9^ per transcribed nucleotide (Methods)^31^. Synthesizing a non-beneficial mRNA of length *l*_*m*_ leads to a growth rate penalty Δ *f*_*m*_ that is linear with the transcription rate^31^, Δ *f*_*m*_ = *c*_*m*_*l*_*m*_*β*_*β*_. For a typical mRNA of length *l*_*m*_ = 1300 nucleotides transcribed at a rate *β*_*m*_ = 30 mRNA / h, the fitness cost of transcription is thus *c*_*m*_*β*_*m*_*l*_*m*_ *≃* 4 *×* 10^−5^ / h (Fig. 3A, Methods), which is selectable^31,82^. The cost of mRNA is also selectable in *E. coli*^24,44^.

**Figure 3.**
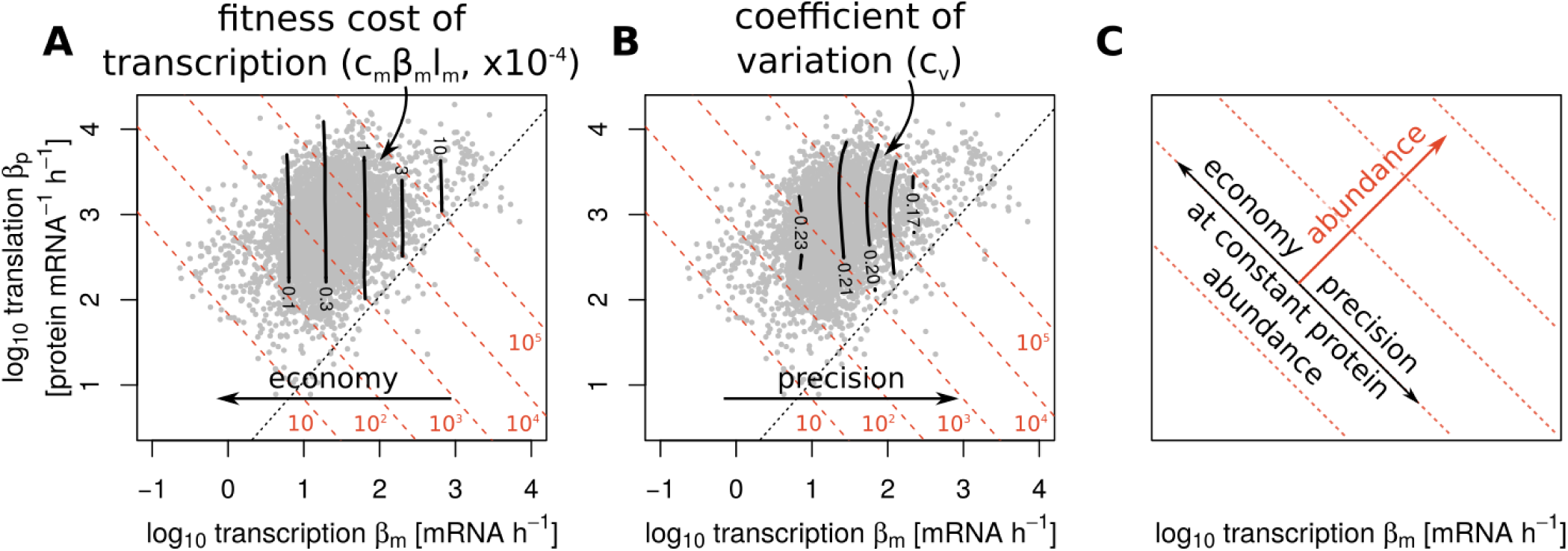
Increasing transcription at constant protein abundance increases transcription cost while decreasing stochastic fluctuations in protein abundance. *S. cerevisiae* rates from Weinberg et al. ^84^. Diagonal dotted lines are lines of constant protein abundance from 10 to 10^5^ proteins per cell. **A.** The loss of fitness *c*_*m*_*β*_*m*_*l*_*m*_ due to transcription (black lines) is linear in the transcription rates *β*_*m*_ and the mRNA length *l*_*m*_. The linear factor *c*_*m*_ (introduced in the next section) rescales transcription fluxes [nt h^−1^] into fitness loss [h^−1^]. In *S. cerevisiae, l_m_* = 1300 nt, *c*_*m*_ = 10^−9^ nt^−1^. **B.** Coefficients of variation (CV, black lines) scale with transcription rates. We applied 2D Gaussian smoothing on CVs (Methods). *S. cerevisiae* data: CVs from Newman et al. ^50^, ribosome profiling and mRNAseq data from Weinberg et al. ^84^. C. Precision in gene expression increases with transcription whereas protein abundance depends both on transcription and translation. Thus, at a given protein abundance, increasing transcription increases the precision of gene expression at the expense of higher transcription costs. See also Figure S3.

In addition to their cost, high transcription rates also have benefits in reducing the noise^5,45,52,54^. Increasing the transcription rate while keeping protein abundance fixed should therefore decrease stochastic fluctuations in protein abundance.

To test if this prediction holds genome-wide across the diversity of chromosomal context and promoters, we use measurements of cell-to-cell variations in protein abundance in S. *cerevisiae*^50^and *E. coli*^76^. Cell-to-cell variations in protein abundance can be quantified by the coefficient of variation (CV). We determine contours of the CV as a function of transcription and translation rate using Gaussian smoothing (Methods), and compare these to contours of protein abundance. Both in S. *cerevisiae* (Fig 3B) and *E. coli* (Fig. S3A), the CV decreases with increasing transcription and decreasing translation on each equi-protein line. The CV mainly scales with transcription, as predicted by theory^54^ (Methods).

Hence, transcription and translation rates impact both gene expression precision and mRNA economy. At a given protein abundance, high translation / transcription ratios lead to economy but also higher gene expression noise, whereas low translation / transcription ratios yield high precision at the expense of higher mRNA cost (Fig 3C). The lack of genes combining high transcription and low translation could be explained by this trade-off: for genes located in the depleted region, the benefits of increased precision may be smaller than transcriptional costs.

### The precision-economy trade-off and the noise floor explain the depleted region of the Crick space

To quantitatively test whether a trade-off between precision and economy can explain the depleted region, we developed a minimal mathematical model of the fitness cost and benefit of transcription and translation (Fig. 4). The model has two main predictions: first that the optimal ratio between translation and transcription rates *β*_*p*_/*β*_*m*_ is set by the ratio of transcription cost per mRNA molecule *C* and the gene’s sensitivity to noise *Q* (defined below). The second prediction is an analytical formula for the boundary of the depleted region — the lower bound *k* on the *β*_*p*_/*β*_*m*_ ratio — based on fundamental parameters.

**Figure 4.**
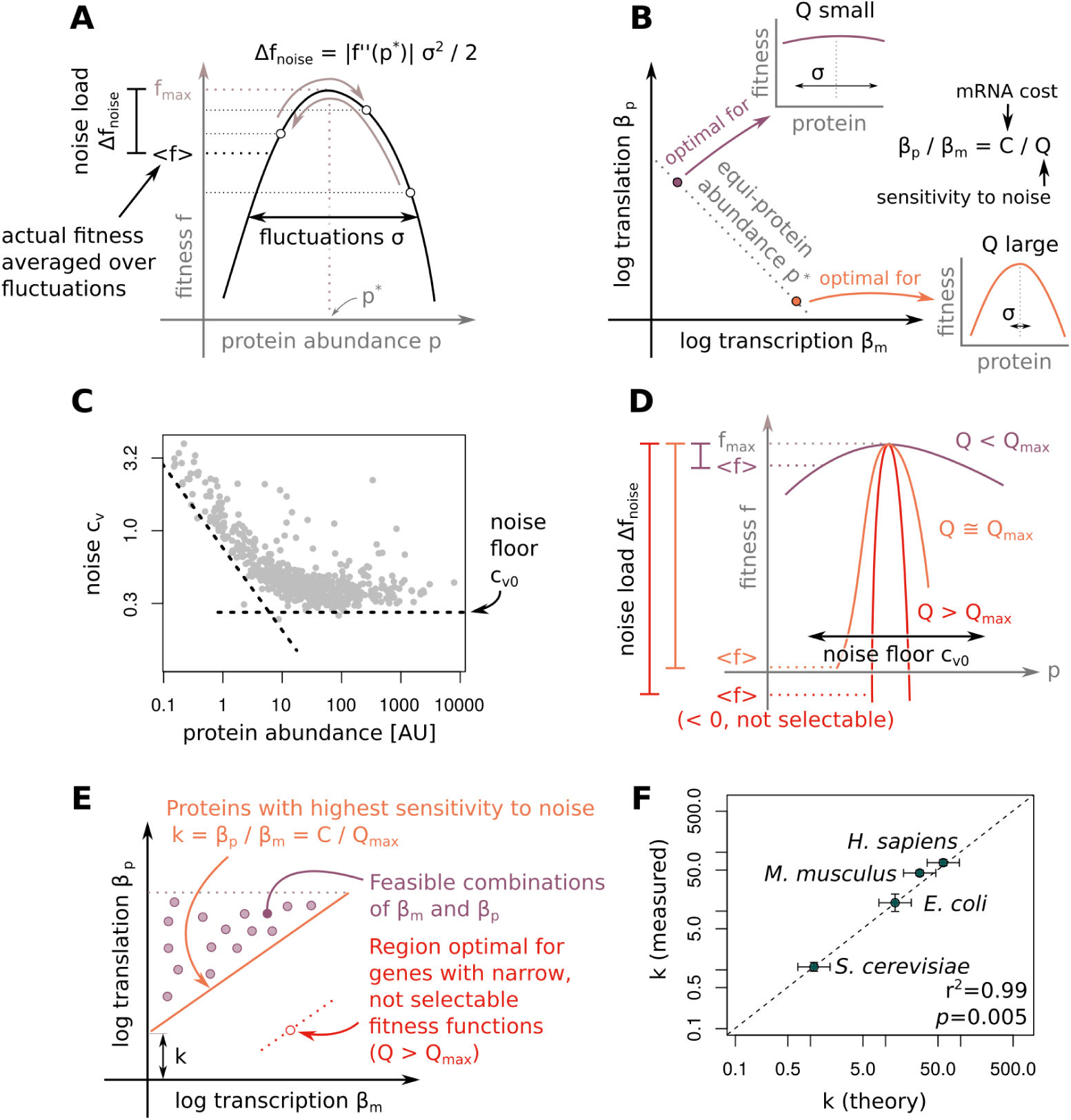
A trade-off between precision and economy explains the depletion of gene combining high transcription with low translation. **A.** The noise load Δ *f*_*noise*_ is the loss of fitness due to stochastic fluctuations in protein abundance. **B.** The optimal *β*_*p*_/*β*_*m*_ depends on the transcription cost per mRNA (*C*) and on how noise sensitive (*Q* ∼ *|f ^″^* (*p\**)*|*) the gene is. Noise sensitive genes have narrower fitness functions (*|f ^″^* (*p\**)*|* large). For a given *p\**, higher transcription *β*_*m*_ decreases fluctuations in protein abundance. Genes that are sensitive to noise (*Q* large) should thus have low translation / transcription ratio. On the other hand, precision is less critical for genes with flat fitness functions (*Q* small). These genes should have higher *β*_*p*_/*β*_*m*_ to keep transcription costs low. **C.** The precision of gene expression is limited by the noise floor *c*_*v*0_. Protein abundance and CV data re-plotted from Taniguchi et al. ^76^. The noise floor is also found in the *E. coli* measurements of Silander et al. ^71^ (Fig. S4B). **D.** Genes with fitness functions narrower than the noise floor *c*_*v*0_ have negative average fitness (*< f ><* 0). Genes with negative fitness are not selectable. Thus, the noise floor *c*_*v*0_ prevents the selection of narrow, noise sensitive fitness functions. **E.** The maximal, selectable noise sensitivity *Q*_*max*_ determines the position *k* of the boundary of the depleted region. Proteins lying on the boundary have maximal noise sensitivity *Q* = *Q*_*max*_. Feasible combinations of transcription and translation correspond to genes that are less sensitive to noise *Q < Q_max_*. **F.** The precision - economy trade-off theory predicts the position *k* of the depleted region in organisms from bacteria to mammals. Error bars represent 95% confidence intervals. See also Figure S4.

To determine the optimal *β*_*p*_/*β*_*m*_ ratio under the precision - economy trade-off, we first model how the *β*_*p*_/*β*_*m*_ ratio affects mRNA economy and precision. We then model how mRNA economy and precision affect fitness. Finally, we determine an analytical expression for the optimal *β*_*p*_/*β*_*m*_ ratio.

To compute how the *β*_*p*_/*β*_*m*_ ratio affects economy, we model the fitness cost of transcription^31^ by the linear function Δ *f*_*m*_ = *c*_*m*_*l*_*m*_*β*_*β*_ where *l*_*m*_ is the (pre-)mRNA length and *c*_*m*_ is the fitness penalty per transcribed nucleotide. *c*_*m*_ can be estimated from the growth rate *µ* and the total transcriptional output Σ *β*_*m*_ if we assume that non-beneficial mRNA are transcribed at the expense of beneficial mRNAs. If a total of Σ *β*_*m*_*l*_*m*_ nucleotides are transcribed in a cell, the average fitness contribution of each nucleotide is *µ/* Σ *β*_*m*_*l*_*m*_. This is also the fitness lost per nucleotide of transcribing a non-beneficial mRNA at the expense of a beneficial mRNA. Thus, the fitness cost per transcribed nucleotide is

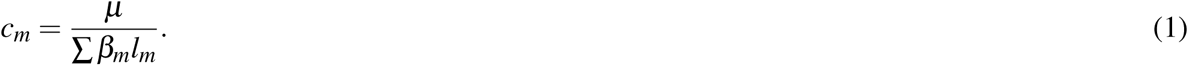

This provides *c*_*m*_ *∼* 10^−9^ in *S. cerevisiae* which agrees well with experimental measurements mentioned above (Methods).

To see how *β*_*p*_/*β*_*m*_ and precision affect fitness, we consider a protein of abundance *p*. The protein contributes a quantity *f* (*p*) to the organism’s fitness (Fig. 4A). Because protein abundance fluctuates around the average expression *< p >*= *p\**, the cell doesn’t experience the maximum fitness *f*_*max*_ but rather a lower average fitness *< f* (*p*) *>*. The fitness lost due to stochastic fluctuations in protein abundance Δ *f*_*noise*_ is called the noise load^32,81^ (Fig. 4A). By expanding the fitness function *f* to second order and averaging over fluctuations in protein abundance, we can compute the noise load:

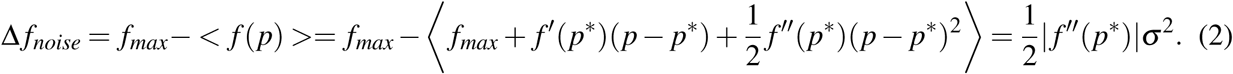

Noise load increases with the curvature of the fitness functions (large *| f ^″^*(*p\**)*|*) and with noise (large *σ*).

To find how *β*_*m*_ and *β*_*p*_ affect fitness through the precision of gene expression, we note that *β*_*m*_ and *β*_*p*_ affect the noise level *σ* ^2^ in a well-characterized way. Theory and experiments^5,50,54,76^ indicate that the variance of protein abundance is given by

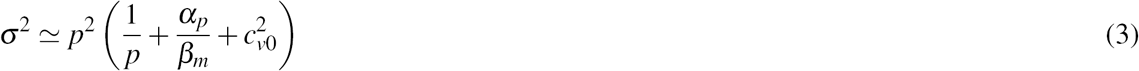

where *α*_*p*_ is the protein decay rate and *c*_*v*0_ is the noise floor (Methods). The noise floor is the minimal amount of cell-to-cell variation in protein abundance in clonal populations^13,50,71,76^.

We can now determine the optimal transcription and translation rates *β*_*m*_ and *β*_*p*_ that minimize the combination of the transcription cost Δ *f*_*m*_ and of the noise load Δ *f*_*noise*_ (Methods),

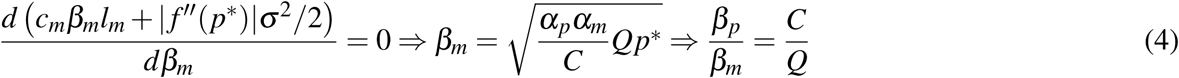

where *C* = *c*_*m*_*l*_*m*_*α*_*α*_ = *c*_*m*_*l*_*m*_*β*_*m*_/*m* quantifies the cost of transcription per mRNA molecule, and 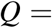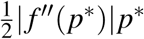 is the gene’s sensitivity to noise. Genes with narrower fitness functions (larger *| f ^″^* (*p\**)*|*) are more sensitive to noise because the noise kicks protein abundance farther from optimum. The sensitivity of a gene to noise also depends on protein abundance *p\** because the noise generally scales with protein abundance *σ* ^2^ *∼ p\**.

Although the cost of translation typically dominates transcriptional cost^31,44,82^, the *β*_*p*_/*β*_*m*_ ratio does not depend on translation cost. This is because translation costs are determined by the total amount of protein translated, regardless of whether the protein is synthesized from few or many mRNAs.

The model therefore predicts a relationship between *β*_*p*_/*β*_*m*_ ratios and the shape of fitness functions (Fig. 4B). Broad fitness functions should have large *β*_*p*_/*β*_*m*_ because genes with broad fitness functions are not sensitive to noise. For those, high precision provides little benefit, and it is best to maximize economy by lowering transcription. On the other hand, genes with narrow fitness functions are sensitive to noise. For those, requirements of high precision to keep the noise load low dominate the cost of transcription. Genes with narrow fitness functions should therefore have small *β*_*p*_/*β*_*m*_ ratios.

We compared the model prediction for the curvature of the fitness function near its peak to recent measurement of fitness functions of 21 genes in *S. cerevisiae*^34^. We find that the curvatures predicted from *β*_*p*_/*β*_*m*_ are within an order of magnitude of the measurement curvatures without any fitting parameters (Fig. S4A). Predictions and measurements correlate positively (*r* = 0.39). A shuffling test suggests that the agreement between the predictions and measurements is unlikely due to chance (*p* = 0.04, Methods). However, variability in the experimental measurements precludes a conclusive comparison.

To find a lower bound *k* on *β*_*p*_/*β*_*m*_ and explain the boundary of the depleted region, we note that there is a limit to how small the noise in gene expression can be. This limit, called the ‘noise floor’ *c*_*v*0_, is revealed by measurements of cell-to-cell variation of protein abundance in clonal populations^13,50,71,76^ (Fig. 4C, Fig. S4B–C). The cause for the noise floor is a current research topic^60^ and it has been proposed that it is due to extrinsic noise^76^ or larger transcriptional burst size of high abundance proteins^13^. The noise floor puts an upper bound *Q*_*max*_ on how noise-sensitive genes can be: for genes with fitness functions narrower than this limit *Q > Q_max_*, the noise load dominates the benefit of expressing the gene (Fig. 4D), leading to negative fitness. Because genes with *Q > Q_max_* cannot be selected for, all endogenous genes must satisfy *Q < Q_max_*. The boundary of the depleted region *β*_*p*_/*β*_*m*_ = *k* hence corresponds to genes with highest sensitivity to noise *Q*_*max*_ (Fig. 4E). The depleted region *β*_*p*_/*β*_*m*_ *< k* corresponds to transcription and translation rates that are optimal for genes with fitness functions too narrow given the noise floor.

To find *k*, we substitute *Q* = *Q*_*max*_ in Eq. 4 for the optimal *β*_*p*_/*β*_*m*_ ratio and rewrite *k* in terms of the noise floor and other cell biology constants (Methods):

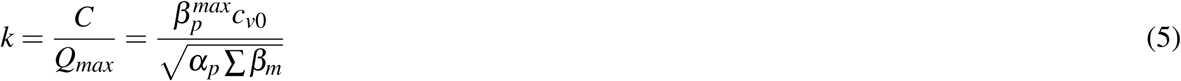

where Σ *β*_*m*_ is the combined transcriptional output of all genes. This expression provides an intuition for the cellular parameters which set the boundary of the depleted region of the Crick space. For example, a larger noise floor *c*_*v*0_ raises the boundary because there is less benefit in having high transcription. An increased transcriptional output Σ *β*_*m*_ lowers the boundary because individual mRNAs are less costly, allowing increased precision in gene expression.

This formula for *k* accurately predicts the boundary of the depleted region of the Crick space (Fig. 4F; red lines on Fig. 2) despite the fact that *k* varies by nearly two orders of magnitude between organisms (Table 1). Thus, the depleted region can be explained in terms of fundamental parameters such as the noise floor, maximal translation rate, total transcription output and mean protein decay rate. Individual cellular constants alone cannot accurately predict *k* (Fig. S4D). Neither can *k* be predicted by noise alone without considering economy, such as hypothesizing that the depleted region is made of all *β*_*m*_ and *β*_*p*_ for which increasing transcription provides little extra precision relative to the the noise floor (Methods).

## Discussion

We find that the distribution of genes in Crick space is not random: genes combining high transcription and low translation are depleted. Such combinations of high transcription and low translation can be achieved with synthetic gene constructs^37^. Therefore, mechanistic constraints cannot explain this depletion. We explain the depletion by a trade-off between precision and economy: increasing transcription at constant protein abundance diminishes stochastic fluctuations, but at a fitness penalty due to the cost of transcription. High transcription rates are therefore optimal for genes that are sensitive to noise whereas low transcription rates are well suited for genes that can tolerate high noise (Fig. 5A). A quantitative model of this trade-off predicts the curvature of the fitness function for each gene and quantitatively explains the boundary of the depleted region.

**Figure 5.**
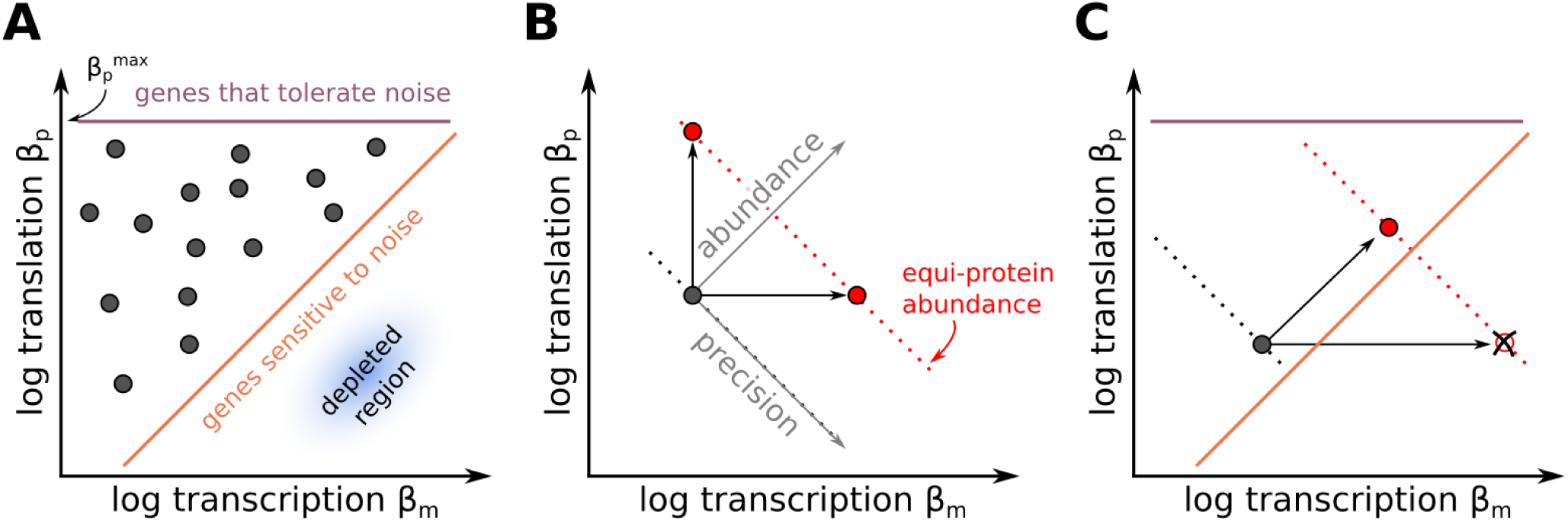
The precision - economy trade-off suggests rules for choosing transcription and translation rates and for selecting regulatory strategies. **A.** Due to the precision - economy trade-off, low *β*_*p*_/*β*_*m*_ is preferred for genes that are sensitive to noise. In log-log scale, low *β*_*p*_/*β*_*m*_ corresponds to a diagonal line. On the other hand, high *β*_*p*_/*β*_*m*_ is preferred for proteins that tolerate noise. Because translation rates have an upper limit 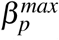, genes with highest *β*_*p*_/*β*_*m*_ are found on a horizontal line. **B.** Regulatory strategies that lead to the same protein abundance differ in how they impact precision. Up-regulating transcription simultaneously increases protein abundance and precision. On the other hand, up-regulating translation increases protein abundance while decreasing precision. Transcription control is thus advantageous assuming that precision is desirable. **C.** When protein abundance changes by a large amount, pure transcription regulation can put the gene in the sub-optimal, depleted region. This can be avoided by co-regulating transcription and translation. See also Figure S5.

The predicted depleted region seems to account for about 99% of the genes. The remaining 1% fall in the depleted region. For some of these genes, evidence from focused studies indicates low translation rates and/or translation control, such as the *E. coli* outliers rimM and trmD^85^, HAC1 in *S. cerevisiae*^61^ and Fth1 in *M. musculus* and *H. sapiens*^58^. *One explanation for such outlier genes is that they face constraints beyond the precision-economy tradeoff. For example, the amino acid response regulator GCN4 in S. cerevisiae* is strongly transcribed and poorly translated in rich medium. But under amino-acid imbalance conditions, a general decrease in protein translation triggers derepression of GCN4 translation through a translation reinitiation mechanism involving short upstream open reading frames^29^. This regulatory mechanism couples GCN4 synthesis to translation stress. It also bypasses transcription, which could allow for rapid upregulation. Such considerations of regulatory couplings or speed might overshadow precision-economy-based limits for certain genes.

The distribution of genes in Crick space is bounded above by the maximal translation rate (10^3.6^ – 10^4^ proteins h^−1^), and below by the boundary of the depleted region. The position of each gene in this space is determined, in the present picture, by the curvature of its fitness function. Genes with a narrow fitness function are most sensitive to noise, and are predicted to lie near the boundary of the depleted region. Genes with a broader fitness function are predicted to lie farther above this boundary. This prediction suggests an experimental test by measuring the curvature of the fitness function and comparing to the prediction. Accurate measurements of fitness functions can be performed by titrating protein concentration experimentally and measuring fitness. A recent experiment measured such fitness functions for 85 genes in *S. cerevisiae*^34^. While we found a statistical agreement between the predicted and measured curvatures of fitness functions, the measurement errors were too large to permit a meaningful comparison to the present predictions (Fig. S4A). Further experiments to measure fitness functions can test whether transcription and translation rates predict fitness curvature near the peak.

Beyond optimizing transcription and translation under a precision-economy trade-off, an additional reason why genes may lie farther above the boundary of the depleted region is the possibility that noise is beneficial for some genes. This occurs for example in cases of bet hedging where a gene product brings little or no fitness advantage at present, but expression is maintained in case conditions change so that the gene product becomes important. In such cases, theory and experiments have shown that a wide cell-cell variation in protein level can be beneficial^19,38,69,87^. Such genes expressed for possible future needs are predicted to lie far above the boundary of the depleted region. This prediction is in agreement with the finding of stress genes at relatively high positions above the boundary, and of essential genes closer to the boundary (Fig. S5A–B, Table S2, Supplementary Item 1).

The theory has further testable experimental predictions. By expressing a protein from different synthetically produced combinations of translation and transcription rates, one should find that there is an optimal translation / transcription ratio. Upon changing growth conditions such that the protein becomes even more important for growth, the optimal transcription rate should increase whereas the optimal translation rate should show little change.

The finding that essential, high-precision genes are located close but not below the boundary of the depleted region has implications for synthetic circuit design. If a protein needs to have a specific abundance for the circuit to functions properly, that protein should be expressed using a promoter and ribosomal binding site (RBS) that puts it close to the boundary of the depleted region. In other words, the optimal design should have *β*_*p*_/*β*_*m*_ = *k* with a value of *k* appropriate to the organism (Table 1). On the other hand, if the circuit is insensitive to the exact concentration of that protein, the protein should be expressed using a weaker promoter and stronger RBS to save transcriptional resources. Combinations of strong transcription (promoters, enhancers) with weak translation (RBS and so on) should be avoided because they incur high transcriptional cost with no extra precision benefit.

The present approach can also help interpret the mode of regulation when the abundance of a protein needs to change. Increasing a protein level can be done by increasing transcription, translation or both. Studies in several organisms indicate that transcription regulation is more prevalent and strong than translation regulation for most genes^15,30,41–43^. The present theory provides a possible explanation for this observation (Fig. 5B). Transcription regulation increases protein abundance and at the same time decreases noise. Translation regulation will increase noise. Thus transcription control is advantageous assuming that precision is desirable. The relatively rare cases of strong translation regulation may be due to considerations of faster response time, or to cases where it is beneficial to reduce precision, such as in bet hedging^38^. One interesting case is when proteins need to be up-regulated from a very low to a very high level. Geometric considerations rule out a purely transcriptional regulation, because this will put the gene into the depleted region; instead, a combined transcription and translation up-regulation is predicted (Fig. 5C).

The present findings suggest that translation / transcription ratios are determined to some extent by rules, such as precision - economy trade-offs. Rules have been proposed in the past to explain features of the complexity of biological systems^62,68,83^. Patterns such as empty regions of the Crick space can help define the rules, similar to the way empty regions of morphospaces in animal morphology can be used to infer potential evolutionary tasks and trade-offs^28,70,77^. It would be interesting to discover additional rules for gene expression in order to interpret the evolved design of cells and to improve engineering of synthetic circuits.

## Acknowledgments

We thank Benjamin Towbin, Shalev Itzkovitz, Ron Milo, Leeat Keren, Naama Barkai, Moshe Kafri, Mihaela Zavolan, Luca Ciandrini, Nir Friedman, Yoav Voichek, Tzachi Pilpel, Idan Frumkin, Dvir Schirman, Noam Stern-Ginossar, Michal Shreberk, Dan Davidi, Daniel Goodman, Tabitha Bucher and members of the Alon lab for scientific discussions and feedback on the manuscript.

This work was supported by the European Research Council under the European Union’s Seventh Framework Program/ERC Grant agreement 249919, and the Israel Science Foundation. UA is the incumbent of the Abisch-Frenkel Professorial Chair. J.H. acknowledges the support of EMBO (ALTF 1160-2012), the Swiss National Science Foundation (P300P3 158472) and the Swiss Society of Friends of the Weizmann Institute.

## Author contributions

Conceptualization, J.H. and U.A.; Methodology, J.H., A.M. and U.A.; Formal Analysis, J.H.; Writing, J.H. and U.A.; Funding Acquisition, J.H. and U.A.

## Methods

### Supplementary discussion

This study is based on an optimality approach to evolutionary biology^53^ whose aim is to explain adaptations found in living organisms in terms of selective forces. This approach has been used to explain, for example, bacterial growth laws^67,79^ or energy landscape in molecular recognition^63^. It is a fruitful approach in the sense that it can suggest new experiments (see discussion).

The quantitative model rests on the assumption that transcription and translation rates can be tuned independently. Although coupling is seen in bacteria^55^ and eukaryotes^26^, that coupling has itself evolved rather than being an absolute constraint. Studies on synthetic promoters and ribosomal binding sites indicate that transcription and translation rates can be changed independently over a wide range^37^.

Another assumption of the theory is that cells are well-adapted to the conditions in which transcription and translation were measured, namely rapid growth in rich medium. Rapid growth is thought to be a key fitness component of microorganisms like *E. coli* and *S. cerevisiae*^51,86^. Rapid growth also occurs in several contexts in mammals, including immune expansion, cancer, development and stem cells in tissues with rapid turn-over. The mammalian HeLa and 3T3 cell lines studied here have likely been selected for fast growth. HeLa cells were collected from cervical cancer, a condition which selects for mutations that provide a growth advantage. Following collection, HeLa cells underwent serial dilutions for 4 months^64^, a procedure which selects for growth^86^. In the case of 3T3 cells, embryonic fibroblasts were collected and underwent serial dilutions for 3-4 weeks^78^. In the process, cell growth first collapses due to cell senescence, and then recovers to levels of freshly collected embryonic fibroblasts, presumably because immortalized mutants take over the population. Hence, both HeLa and 3T3 cells are likely well adapted to the conditions in which transcription and translation were measured, namely rapid growth in rich culture medium.

### Data sources and estimation of cellular constants in the four model organisms

#### S. cerevisiae

We obtained the processed Reads Per Kilobase per Million (RPKMs) of the mRNAseq and ribosome profiling experiments of Weinberg et al. ^84^ from GEO (GSE75897). We used the RiboZero mRNAseq experiment. Experimental measurements estimate *N*_*m*_ = 60000 mRNA copies per cell^88^ cellular volume at 37*µm*^3^ (BNID100430^48^). Given a protein concentration^46^ of 3 *×* 10^6^*/µm*^3^, we estimate that there are *N*_*p*_ *≃* 1.1 *×* 10^8^ proteins per cell. We used a cell division time is 99min (BNID101310). Given that the median protein half-life is 45min^7^, the typical protein decay rate is *α*_*p*_ = 60 (log(2)*/*45 + log(2)*/*99) = 1.34*h*^−1^. Eser et al. ^18^ estimated the typical mRNA decay rate at *α*_*m*_ = 5.1*h*^−1^. Multiple experiments found a noise floor *c*_*v*0_ *≃* 0.1^5,33,50^.

#### E. coli

From the sequence reads archive, we downloaded the mRNAseq reads and ribosome profiling reads from the experiments Li et al. ^41^ performed in rich medium (mRNAseq: SRR1067773, SRR1067774, ribosome profiling: SRR1067765, SRR1067766, SRR1067767, SRR1067768). We obtained the *E. coli* genome sequence and transcriptome annotation from NCBI (accession NC 000913.3).

All genome mappings were performed using Bowtie2in local alignment mode. We discarded all technical reads as well as reads that mapped against non-coding RNAs, defined as transcripts marked as ‘ncRNA’, ‘rRNA’ or ‘tRNA’ in the genome annotation. Remaining reads were mapped to transcripts marked as ‘CDS’ in the genome annotation. Ribosome profiling reads were mapped to coding transcripts after trimming the first and last 5 codons to remove the effect of translation initiation and termination. Reads that mapped equally well to multiple loci were assigned to one of the loci at random. We then computed RPKMs per gene. Because reproducibility between runs was high, we combined reads from all runs for subsequent analyses.

The resulting mRNA abundances and protein synthesis rates estimates were highly correlated with those computed by Li et al. ^41^ (*r*^2^ *>* 0.99). Differences could be due to the updated genome version we used (NC 000913.3 vs NC 000913.2 in the original analysis of Li et al.), differences in the aligner (we used Bowtie2 while Li et al. ^41^ used Bowtie), and other differences in the implementation of the bioinformatics pipeline. Repeating all the *E. coli* analyses using the protein synthesis rates from Supplementary Table S1 and mRNA abundance Supplementary Table S4 of Li et al. ^41^ leads to minor changes in the exact position of genes in Crick space and supports all conclusions presented in this article.

There are 1380 mRNAs per cell (BNID100064). Cell volume is *≃* 1*µm*^3^ (BNID100004). Assuming a protein concentration^46^ of 3 *×* 10^6^*/µm*^3^, there are about 3 *×* 10^6^ proteins per cell. Doubling time was measured by Li et al. ^41^ at 21.5 minutes, which puts the growth rate at *µ* = 60 log(2)*/*21.5 = 1.93*h*^−1^. Since ribosome profiling RPKMs correlate well with protein abundance^41^, we neglect protein degradation (*α*_*deg*_ = 0, *α*_*p*_ = *µ*). The median mRNA half-life is 2.8 minutes^11^, which corresponds to decay rate *α*_*m*_ = 14.9*h*^−1^. The noise floor *c*_*v*0_ was measured to be about 0.25: 0.27 *±* 0.01 in Taniguchi et al. ^76^, 0.22 *±* 0.01 in Silander et al. ^71^.

#### M. musculus

We downloaded the processed RPKMs of the mRNAseq and ribosome profiling experiments of^16^ from GEO (GSE60426).

Given a 3T3 cell volume of *V* = 2000*µm*^3^ ^66^ and a protein concentration of 3 *×* 10^6^ *µm*^−3^ ^46^, there are *N*_*p*_ *≃* 6.0 *×* 10^9^ proteins per cell. Measurements suggest that there are around *N*_*m*_ = 180000 mR-NAs per 3T3 cell^66^. Following experimental measurements^66^, we used a cell cycle time of 24h and a protein half-life of 48h (after removing the effect of cell division^66^). These numbers correspond to a growth rate *µ* = 0.03*h*^−1^ and a degradation rate *α*_*deg*_ = 0.01, which puts the protein decay rate at *α*_*p*_ = *µ* + *α*_*deg*_ = 0.04*h*^−1^. Friedel et al. ^21^ found a median mRNA decay rate of *α*_*m*_ = 0.14*h*^−1^ in 3T3 cells, which is the value we used here. Another study^66^, also in 3T3 cells, measured a median half-life of *α*_*m*_ = 0.08*h*^−1^, a value for which the predicted boundary is also in good agreement with rates measurements (Fig. S2I). To our knowledge, the noise floor *c*_*v*0_ hasn’t been measured in mouse. We therefore used the noise floor from the closest organism in evolutionary terms, namely *H. sapiens*.

#### H. sapiens

We downloaded the processed RPKMs of the mRNAseq and ribosome profiling experiments of^16^ from GEO (GSE60426).

Given a cellular volume of *V* = 2500*µm*^3^ (BNID103725) and a protein concentration^46^ of 3 *×* 10^6^*/µm*^3^, there are about *N*_*p*_ = 7.5 *×* 10^9^ proteins per cell. A division time of 22h (BNID109393) corresponds to a growth rate *µ* = 0.03*h*^−1^. The protein degradation rate measurements of Cambridge et al. ^9^ found *α*_*deg*_ = 0.02. This puts the effective protein decay rate at *α*_*p*_ = *µ* + *α*_*deg*_ = 0.05*h*^−1^. We could not find direct measurements of the number of mRNAs per HeLa cell *N*_*m*_. A back of the envelope calculation puts *N*_*m*_ in the 10^5^ – 10^6^ range^47^. This is consistent with recent smFISH and RNAseq measurements which found 10^5^ mRNAs in MIN6 and 10^6^ mRNAs in liver cells^4^. Starting with the 180000 mRNAs per 3T3 cells^66^ and assuming that mRNA content scales with cell volume, we estimate *N*_*m*_ *≃* 180000 *×* 2500*/*2000 = 225000 mRNA per HeLa cell. The accuracy of the predicted boundary of the depleted region is robust to halving or doubling *N*_*m*_ (112500 *< N_m_ <* 450000, see Fig. S2J–K). Gregersen et al. ^25^ measured mRNA half-lives in (human) HEK293 cells and found a median half-life of 11.4h, which is the value we used here (*α*_*m*_ = 0.06*h*^−1^). Another study measured a median half-life of 5h in human B-cells (BL41)^21^, a value for which the predicted boundary of the depleted region is also in good agreement with experimental data (Fig. 2L). Dar et al. ^13^ found a noise floor *c*_*v*0_ *≃* 0.3.

### Estimating transcription and translation rates from high-throughput experiments

For each gene *i*, we estimated the number of mRNAs per cell *m*_*i*_ from the total number of mRNAs per cell *N*_*m*_ from the per-gene mRNAseq RPKM *r*_*i*_ data as

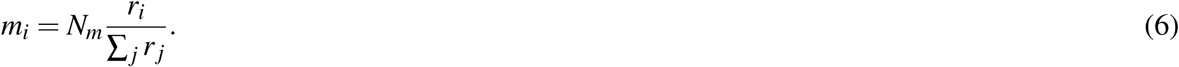

At steady-state, mRNA abundance *m* is the ratio of the transcription rate *β*_*m*_ to the mRNA decay rate *α*_*m*_ ^27^. We thus estimated the transcription rates of each gene as

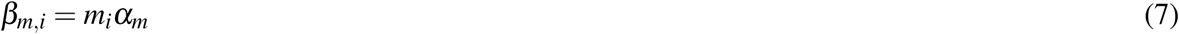

where *α*_*m*_ is the median mRNA decay rate (Table S1). Finally, we estimated translation rates *β*_*p*_ by combining three numbers: the total number of proteins per cell *N*_*p*_, the protein decay rate *α*_*p*_, and the gene’s ribosome profiling RPKM *s*_*i*_ of gene *i*. The number of proteins synthesized per time unit is *N*_*p*_α_*p*_. A fraction *s*_*i*_/ Σ_*i*_ *s*_*i*_ of this protein synthesis flux is translated from mRNAs *m*_*i*_ of gene *i*. We estimate the translation rate *β*_*p,i*_ (expressed per mRNA copy per cell) by dividing the protein synthesis of each gene by the mRNA copy number *m*_*i*_:

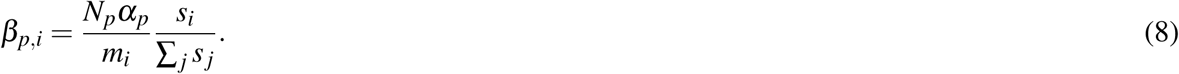

Estimates of *N*_*p*_ and *α*_*p*_ are provided in Table S1.

### Accounting for gene-specific mRNA and protein decay rates has a small impact on the position of genes in 2D Crick space

To evaluate the effect of this simplification, we plot genes in 2D Crick space taking gene-specific mRNA and protein decay rates into account (Fig. S1E). We then assign the same median mRNA and protein decay rates to all genes to re-estimate transcription and translation rates (Fig. S1F). The gene positions in the two resulting 2D Crick spaces differ by less than 0.3 (root mean square deviation in log^10^ rates), which is ≃ 10% of the total variation (about 3 log^10^ units in transcription and translation rates). We conclude that taking into account gene-specific mRNA and protein decay rates has only a small impact on the position of genes in 2D Crick space and thus on present conclusions.

### Sequencing data processing

We consider only genes for which the measurement error was small enough to allow accurate estimation of transcription and translation rates. Accurate estimation of these rates is difficult for low abundance mRNAs because they may only collect a handful of reads. This leads to a large uncertainty on the mRNA copy number *m*, and thus on the transcription rate *β*_*m*_ = *mα_m_*. How many reads per gene should we require to be confident about our estimates of transcription rates?

Estimates of mRNA abundance *m* scale with the number of reads *n* mapping to a given gene (relative to the gene length). We thus compute the minimum number of reads per gene needed to keep the sampling noise on log^10^ mRNA abundance below a certain threshold *ε*

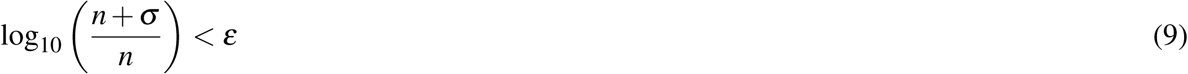

where *σ* is the standard deviation on *n* due to the sampling error. We model sequencing as a Poisson process, and thus 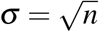. Substituting this expression for *σ* into Eq. 9, we compute the minimal number of reads necessary to control for a given error *ε* on log^10^ mRNA abundances:

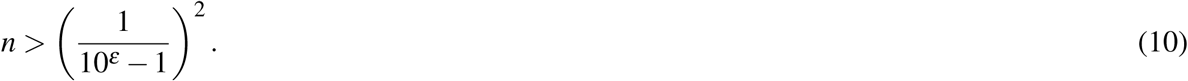

A minimum of 10 reads per mRNA is needed to keep the sampling error on log^10^ transcription rates in the *±*0.1 range (Fig. S2M). We therefore discard genes with less than 10 reads per gene, a procedure which keeps the sampling error low while keeping as many genes as possible in the analysis. Similarly, we require at least ribosomal profiling 20 reads per gene. We applied the same criteria to all four organisms. We repeated the analyses keeping only genes with at least 100 ribosome profiling reads and reached the same conclusions as the one presented in the article.

In *M. musculus* and *H. sapiens*, we discarded canonical histone genes from the analysis because their mRNAs lack a polyA-tail. The polyA+ selection step of mRNAseq discriminates against these mRNAs. Consequently, the abundance of canonical histone mRNAs is underestimated by mRNAseq RPKMs, leading to aberrant (high) translation rate estimates.

### Data and Software Availability

We deposited the raw data (RPKMs from RNAseq and ribosomal profiling) from which we estimated transcription and translation rates at this address: https://data.mendeley.com/datasets/2vbrg3w4p3/draft?a=955cbbdf-9f26-4fbb-970b-e6b4081c1f3e Estimated transcription and translation rates are also found at that address. In addition, downloadable tables contain extra fields (such as coefficient of variation on protein abundance fluctuations) needed to reproduce Fig. 3B and 4C.

### Estimating maximal translation rates

To estimate the maximal translation rate, we ask how fast proteins can be translated 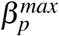 from a single mRNA in the limit where translation initiation is no longer limiting. In this regime, ribosomes follow each other closely along the mRNA. A given ribosome needs to move forward before the next one can advance. The speed at which ribosome elongate the peptide chain and how many codons each ribosome occupies on the mRNA determine how fast proteins can be synthesized.

If a ribosome occupies *L* codons on the mRNA and *v* codons are translated into amino-acids per second, it takes *L/v* seconds for a ribosome to free space for the next ribosome. The maximal flux of ribosomes at given codon is thus *v/L*. This flux sets an upper bound on the translation rate: 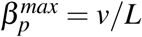.

In *S. cerevisiae*, the elongation rate *v* in favorable growth conditions is 10 amino-acids per second (BNID107871). Each ribosome occupies 28 nucleotides (BNID107874), so L=9.3 codons. This puts the maximal translation rate 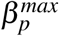 at 10^3.6^ proteins per hour. This bound is in good agreement with estimated translation rate in *S. cerevisiae* and other eukaryotes.

In *E. coli*, translation rate can be up to 10^4^ proteins per hour (Fig. 2). While the size of ribosomal footprints have not been determined, prokaryotes have smaller ribosomes (21nm, BNID102320) than eukaryotes (26.5nm, BNID111542) and so ribosomal footprints should be smaller. Assuming that ribosomal footprints are proportional to the size of the ribosome, we estimate that prokaryotic ribosomes cover 22 nt or *L* = 7.3 codons. In favorable conditions, *E. coli* can elongate up to *v* = 21 amino-acids per second (BNID100059). This leads to a maximal translation rate of 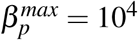 proteins per hour.

### Assessing the statistical depletion of genes combining high transcription with low translation

To test for a statistical significant depletion of genes combining high transcription with low translation, we used a randomization strategy.

First, we defined the depleted region as the sub-region of the Crick space lying below a line of slope 1 that has 1% of the genes below it. We then asked if this figure of 1% was high or low compared to chance.

To find out, we randomized transcription and translation rates. Because the main goal of gene expression is to express proteins at the right abundance, we required that randomized datasets have the same distribution of protein abundance as the original dataset. Also, we enforced the observed upper bound *β*_*p,max*_ on translation rates. To do so, we randomly sampled a protein abundance *p* and a translation rate *β*_*p*_ for each gene. We then computed the corresponding transcription rates *β*_*m*_ = *pα_m_α_p_/*β*_p_*, with *α*_*m*_ and *α*_*p*_ the mRNA and protein decay rates reported in Table S1. Finally, we determined what fraction of genes in the randomized dataset were found below the line of slope 1 and leaving 1% of the genes of the original dataset below it.

We repeated the procedure 10^4^ times to determine the distribution of the fraction of genes in the depleted region expected by chance. We finally estimated the *p*-value that genes avoid combining high transcription with low translation from the fraction of randomized datasets with more genes in the depleted region than the original dataset.

### Estimating transcription and translation rates in the synthetic gene library of Kosuri et al

We compared the distribution of transcription and translation rates of *E. coli* genes to that of the synthetic constructs library of Kosuri et al. ^37^. This study quantified the mRNA and protein abundance of each construct. The constructs only differed in their ribosomal binding sites and promoters. We thus assumed that they shared the same mRNA decay rates and protein decay rates. As a result, the transcription rate of each construct is proportional to mRNA abundance. Translation rates are proportional to the ratio of protein abundance to mRNA abundance.

To compare these measurements to our absolute transcription and translation rates estimates of *E. coli* genes, we assumed that the strongest promoters and RBSs of Kosuri et al. ^37^ yielded transcription and translation rates comparable to *E. coli*’s strongest promoters and RBS. We did so by aligning the 99th percentiles of the transcription and translation rates of the synthetic constructs to those of *E. coli* genes.

The conclusions of the comparison are robust to this assumption. For instance, even if we assume that the strongest Kosuri promoters and RBSs achieve transcription rates 10 times higher or lower than the strongest *E. coli* promoters (i.e. shifting the red cloud of Fig. 2E to the right or to the left by one unit), the high transcription - low translation region would still be covered by a sizable fraction of synthetic constructs.

### Expression for the fitness cost of transcription

Experiments and theory suggest that the fitness cost of transcription Δ *f*_*m*_ scales with the transcription rates^24,31^ and (pre-)mRNA length^10,59^. For a pre-mRNA of length *l*_*m*_ and transcription rate *β*_*m*_, we thus write the fitness cost of transcription Δ *f*_*m*_ as

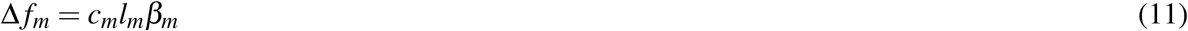

where the constant *c*_*m*_ rescales transcription fluxes (nucleotides per hour) into fitness units (per hour). In this section, we estimate the proportionality constant *c*_*m*_.

To do so, we hypothesize that transcriptional resources are limiting. In this case, making one non-beneficial mRNA comes at a cost because it replaces a fitness-contributing mRNA. The average fitness contribution of a useful mRNA is 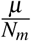 where *µ* is the growth rate and *N*_*m*_ = Σ *β*_*m*_/α_*m*_ is the total number of mRNAs per cell. Therefore, the fitness cost of making *m* = *β*_*m*_/α_*m*_ mRNAs is

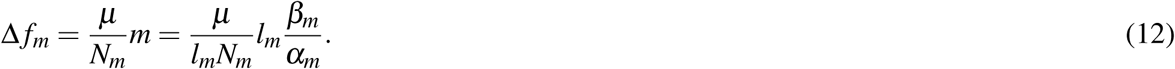

By identifying the terms in Equations 11 and 12, we see that *c*_*m*_ can be estimated from the growth rate *µ*, the typical pre-mRNA length *l*_*m*_ and the total transcriptional capacity Σ *β*_*m*_

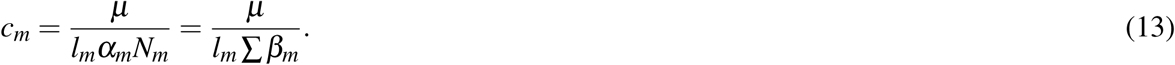

*c*_*m*_ has units of nt^−1^. Alternatively, *c*_*m*_ can also be expressed per mRNA copy (which we will use in the next section):

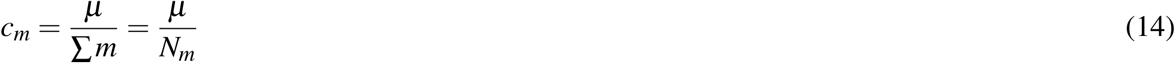

where *N*_*m*_ is the number of mRNAs per cell.

### The fitness cost of synthesizing mRNA in *E. coli* can be predicted from cellular constants

In this section, we show that the expression for *c*_*m*_ derived in the previous section predicts the fitness cost of synthesizing mRNA in *E. coli*. For this, we use the data of Kosuri et al. ^37^ who quantified the abundance of 10,000 different clones that express a fluorescent protein under the control of different promoters and RBSs. As a result, different clones express the fluorescent protein at a different abundance *p*, from a different number of mRNAs *m*.

We model mRNA and protein cost *c*_*m*_ and *c*_*p*_ as linear penalties on the growth rate *µ*

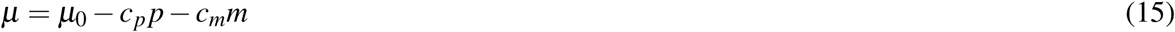

where *µ*_0_ is the rate at which cells would grow in the absence of a fluorescent protein construct. If cells are growing exponentially for *t* hours, the concentration *x*_*m,p*_(*t*) of a clone that expresses *m* non-beneficial mRNAs and *p* proteins is

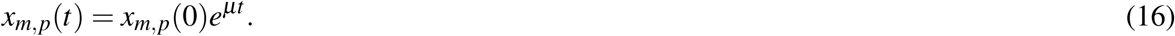

We can normalize *x*_*m,p*_(*t*) to the concentration of clones *x*_0,0_(*t*) that express *m* and *p* at low levels and hence don’t experience a growth penalty (*µ ≃ µ*_0_):

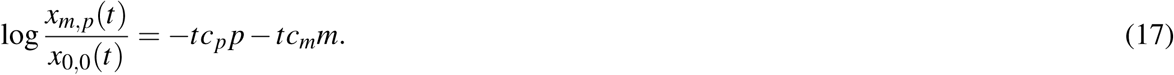

Here, we have assumed that the transformation efficiency of clones is independent of *m* and *p* (*x*_*m,p*_(0) *≃ x*_0,0_(0)). We estimate *x*_*m,p*_(*t*)*/x*_0,0_(*t*) from the ratio between the DNA counts of each clone and the DNA counts of clone that expressed low levels of GFP (prot *<* 1.5 *×* 10^3^ in Table S3 of Kosuri et al. ^37^).

To test if mRNA cost is selectable, we perform two linear regression analyses: one regression of log 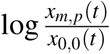 on *p* alone, and one regression of log 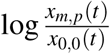 on *p* and *m*. Using the F-test for nested linear models, we find that the squared residuals for the regression on *m* and *p* are significantly smaller than the squared residuals for the regression on *p* alone (*p <* 10^−15^). Therefore, a model that accounts for mRNA and protein cost is significantly more accurate at predicting fitness than a model that account for protein cost alone. This suggests that the cost of synthesizing mRNA is selectable in *E. coli*.

The linear regression estimates of mRNA and protein cost are *tc*_*m*_ = 4.0 *×* 10^−2^ h mRNA^−1^, and *tc*_*p*_ = 2.2 *×* 10^−6^ protein^−1^.

Since the growth time *t* is not known precisely, we cannot determine *c*_*m*_ and *c*_*p*_ individually. But we can determine their ratio: *c*_*m*_/*c*_*p*_ ≃ 630. The fitness cost of transcription theory introduced in the previous section (Eq. 14) predicts

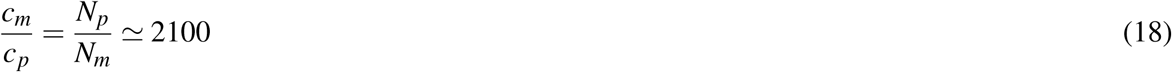

where *N*_*p*_ is the total number of proteins per cell and *N*_*m*_ is the total number of mRNAs per cell (Table S1). Given the typical uncertainty on measurements of *N*_*p*_ and *N*_*m*_ (2-fold), the 95% confidence for *c*_*m*_/*c*_p_ ranges from 300 to 14000. We thus find that predictions of mRNA and protein cost agree with the measurements of Kosuri et al. ^37^.

Finally, we test whether the theoretical estimates for *c*_*m*_ and *c*_*p*_,

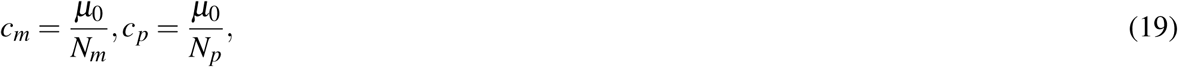

can predict the abundance of clones. While we need to know the growth time *t* to predict clone abundance, the correlation between measured and predicted clone abundance is independent of *t* (see Eq. 17 which relates clone abundance to the growth time and the costs of mRNA and protein).

We find a positive correlation between predictions and measurements of clone abundance (*r* = 0.67, *p <* 10^−15^). We set *t* to one day (*t* = 24h) in Fig. S3B to illustrate the correlation.

In conclusion, the cost of mRNA synthesis in *E. coli* can be predicted from the growth rate and the total transcription output.

### The cost of transcription in *S. cerevisiae* estimated from the measurements of Kafri et al. can be predicted from cellular constants

Here we estimate the growth penalty of transcription in *S. cerevisiae* from the measurements of Kafri et al. ^31^. This study introduced a fluorescent protein construct of pre-mRNA length *l*_*m*_ at different copy numbers *n* in the *S. cerevisiae* genome. The protein abundance *p* of the fluorescent protein depended on the genomic copy number of the construct, as did the transcription rate *β*_*m*_.

This study found that the growth rate *µ* decreases linearly with the genomic copy number *n* of the fluorescent protein construct,

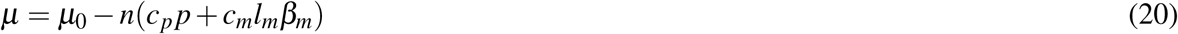

where *µ*_0_ is the growth rate of the WT strain, *c*_*m*_ is the growth penalty of transcription (expressed per transcribed nucleotide) and *c*_*p*_ is the protein burden (growth penalty per protein). To compare cost across growth conditions, the study normalized the growth rate *µ* of strains with genomic insertions of the construct to that of WT *µ*_0_ (e.g. Fig. 4B of Kafri et al. ^31^):

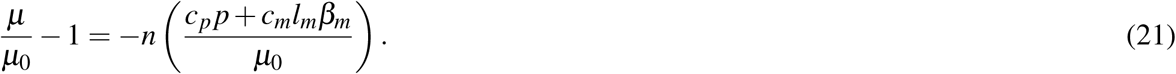

Following the notation of Kafri et al. ^31^, we call *s*_*N*_ the slope of the relative growth rate *µ/µ*_0_ as function of the genomic copy number of the construct *n*. We can write *s*_*N*_ in terms of the fitness cost parameters *c*_*m*_ and *c*_*p*_:

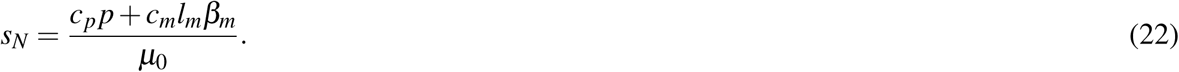

To distinguish between the cost of protein and transcription, Kafri et al. ^31^ designed a second construct (DAmP) lacking a terminator, which makes the mRNA unstable. This reduces protein synthesis between 10 fold (according to protein fluorescence) and 30 fold (according to qPCR) with minimal effect on transcription^31^. Similar to *s*_*N*_, Kafri et al. ^31^ defined *s*_*D*_ as the slope of *µ/µ*_0_ as a function of the genomic copy number *n* of the DAmP construct. We can also write *s*_*D*_ as a function of fitness cost parameters *c*_*p*_ and *c*_*m*_:

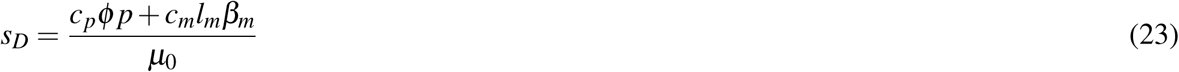

where the factor *ϕ* accounts for the decrease in protein synthesis of the DAmP strains.

From experimental measurements of *s*_*N*_, *s*_*D*_, ϕ, *l*_*m*_ and *β*_*m*_, one can estimate the translation cost per nucleotide *c*_*m*_:

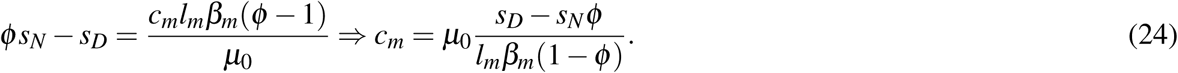

Experiments in YPD medium^31^ measured the following values: *s*_*N*_ *≃* 9.0 *×* 10^−3^, s_*D*_ *≃* 4.1 *×* 10^−3^, *l*_*m*_ *≃* 1000 nt / mRNA, *ϕ ≃* 0.06, *β*_*m*_ *≃* 1300 mRNAs h^−1^, *µ*_0_ *≃* 0.42 h^−1^. Plugging in these values in Eq. 24 estimates the fitness cost *c*_*m*_ per transcribed nucleotide,

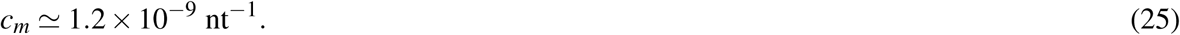

All parameters are known up to two significant digits, except for *ϕ* (10 *≤ ϕ^−1^ ≤* 30), which is known up to one significant digit. This puts the measurement uncertainty at *±*1 *×* 10^−9^ *nt*^−1^.

Plugging the cellular constants of *S. cerevisiae* from Table 1 into the formula for *c*_*m*_ (Eq. 13) estimates the transcription *cost at cm = µ_0_/ Σ *β*_m_l_m_ ∼* 1.2 *×* 10^−9^ nt^−1^. This value is in excellent agreement with the experimental measurements of Kafri et al. ^31^. Such an agreement between experimental fitness parameters in the order of the 10^−9^ may appear surprising given the typical accuracy of biological measurements. The reason for such a good agreement is that the parameters are expressed per transcribed nucleotide whereas actual measurements were performed for full-length mRNAs of a highly transcribed *S. cerevisiae* gene. Because both the transcript length *l*_*m*_ and the transcription rates *β*_*m*_ are in the order of 10^3^ in the experiments of Kafri et al. ^31^, we are actually comparing fitness parameters in the order of 10^−3^, which are accessible experimentally.

### Plotting how the coefficient of variation scales as a function of transcription and translation

We obtained measurements of the coefficients of variation *c*_*v*_ of *S. cerevisiae* and *E. coli* genes from the studies of Newman et al. ^50^ and Taniguchi et al. ^76^. To determine the contours of *c*_*v*_ as a function of transcription and translation, we first applied a Gaussian smoother. The smoother estimates the coefficient of variation *c*_*v*_ for given transcription *β*_*m*_ and translation *β*_*p*_ rates from a weighted average of genes with similar *β*_*m*_ and *β*_*p*_. Genes with comparable *β*_*m*_ and *β*_*p*_ weight more in the average than genes with very different *β*_*m*_ and *β*_*p*_.

Formally, we estimated *c*_*v*_(*β*_*m*_, *β*_*p*_) as a weighted average,

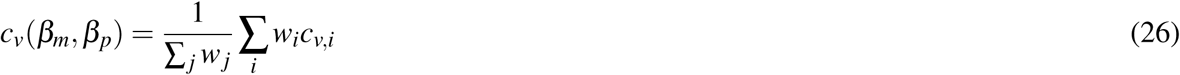

where the Gaussian weights *w*_*i*_ are defined as:

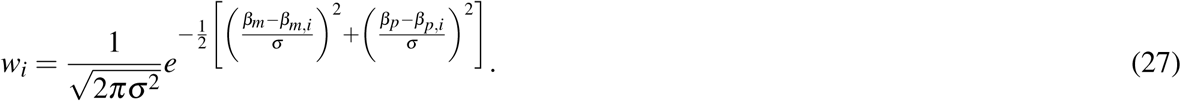

We set the smoothing width *σ* to one fifth of the data range. We only plotted contours of *c*_*v*_(*β*_*m*_, *β*_*p*_) for densely populated regions of the Crick space (Σ *w*_*i*_ *≥* 200).

### Expression for the variance in protein abundance as a function of the Crick rates

In this section, we derive an expression for the coefficient of variation *c*_*v*_ as a function of the transcription rate *β*_*m*_, the protein abundance *p* and the protein decay rate *α*_*p*_. Following a extensive line of theoretical and experimental research^54,72^, we model gene activation and inactivation as a telegraph process (Fig. S3C). Genes are activated at a rate *k*_*on*_ and inactivated at a rate *k*_*off*_. Active genes synthesize mRNAs at a rate *δ*. Messenger RNAs are translated into proteins at a rate *β*_*p*_ and degrade at a rate *α*_*m*_. At steady-state, the coefficient of variation *c*_*v*_ on protein abundance of this stochastic process can be computed analytically^54^:

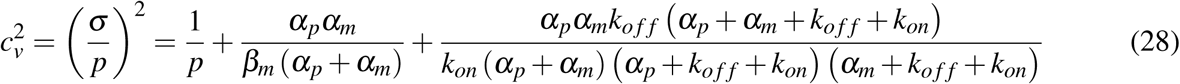

The first term of the equation accounts for the Poisson noise on protein abundance stemming from the protein birth-death process. The second term accounts for the noise caused by translating proteins from mRNAs of low copy number. The last term models the noise caused by gene activation and inactivation and transcriptional bursting.

At present time, it is difficult to measure *k*_*off*_ and *k*_*on*_ genome-wide experimentally. We therefore seek a simplified, approximate expression for the coefficient of variation *c*_*v*_ in which these parameters occur only implicitly through the transcription rate *β*_*m*_. Note that *β*_*m*_ and *k*_*on*_, *k*_*off*_ are related to each other,

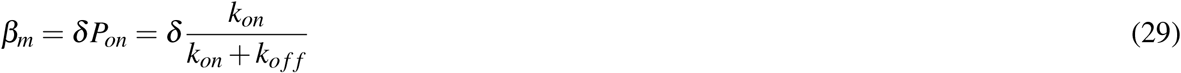

where *δ* is the transcription rate when the gene is in the ‘on’ state, and *P*_*on*_ is the fraction of time when the gene is active.

***E. coli*** With a median half-life of 2.5 min^11^, mRNAs decay much faster than proteins (*α*_*m*_ » α_*p*_). In addition, protein decay *α*_*p*_ is mainly set by the cell division time (20 min or longer)^35^, which is slow compared to gene inactivation which takes place at the time-scale of seconds^72^ (*k*_*off*_ » α_*p*_). In this regime, we can approximate the analytical expression for the coefficient of variation (Eq. 28) as:

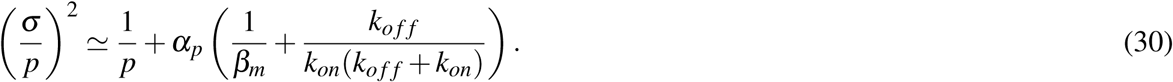

The gene activation rate *k*_*on*_ *≃* 10/h is largely constant across *E. coli* promoters, in contrast to *k*_*off*_ which determines the transcription rate *β*_*m*_ ^72^. Using Equation 29 which relates the transcription rate *β*_*m*_ to gene (in-)activation rates *k*_*on*_ and *k*_*off*_, we can rewrite the coefficient of variation in terms of *β*_*m*_,

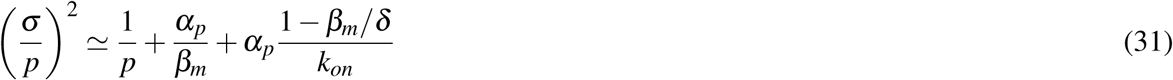

where *δ ∼* 800 / h and *k*_*on*_ *≃*10 / h^72^.

The Poisson noise term 1*/p* is typically negligible compared to the two other terms. The small mRNA copy number noise term *α*_*p*_/*β*_*m*_ dominates at low *β*_*m*_ (Fig. S3D). This term is consistent with the observation that protein noise initially decreases with protein abundance (Fig. 4C, Fig. S4B–C), and that protein noise decreases with transcription (Fig. 3B).

The third term (gene activation noise) becomes dominant for large *β*_*m*_ (Fig. S3D). Because the third term is almost a constant for physiological values of *β*_*m*_ (Fig. S3D), we ask whether it could explain the noise floor found in the single cell experiments of Taniguchi et al. ^76^ and Silander et al. ^71^.

In the growth conditions of Taniguchi et al. ^76^, the doubling time was 150 min which implies a protein decay rate *α*_*p*_ = 0.28/h. Plugging this *α*_*p*_ in Eq. 31, the third term is about 0.03, which is two-fold below the noise floor 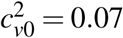 observed in the measurements of Taniguchi et al. ^76^ (Fig. 4C). In Silander et al. ^71^ grew *E. coli* in M9 + 0.2% arabinose, a condition in which *α*_*p*_ *≃*0.45/h^1^. With this *α*_*p*_, the third term is about 0.04, which is comparable to the noise floor 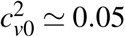 in the measurements of Silander et al. ^71^ (Fig. S4B).

We conclude that gene activation noise may explain the noise floor in the measurements of Silander et al. ^71^. In the experiments of Taniguchi et al. ^76^, gene activation noise is too small to explain the noise floor. There, the noise floor may be explained by other factors such as as extrinsic noise^17,76^. Independently of the specific cause of the noise floor, we model it as a phenomenological constant 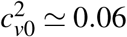 inferred from the measurements of Taniguchi et al. ^76^ and Silander et al. ^71^ (or *c*_*v*0_ *≃* 0.25). This yields an expression for the noise that is accurate with both datasets,

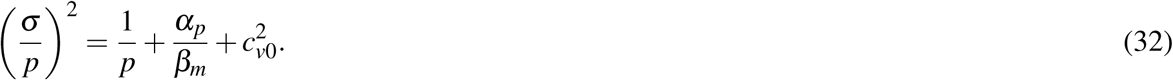

#### S. cerevisiae, H. sapiens, M. musculus

In eukaryotes, an approximate expression for the coefficient of variation can also be derived, but is slightly more complicated because the separation of time-scales is less clear than in *E. coli*: messenger RNAs decay typically faster than proteins, but not by a full order of magnitude. A more realistic, data-driven assumption is

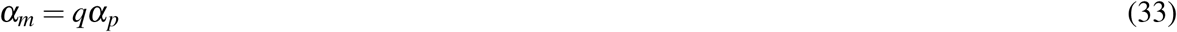

with *q* ≃ 3 (Table S1). Measurements in *S. cerevisiae*^88^ and *H. sapiens*^13^ suggest that gene activation dynamics are much faster than protein decay,

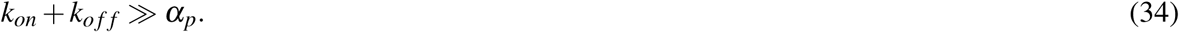

Under these assumptions, we can approximate the analytical expression for the coefficient of variation *(Eq. 28)* as

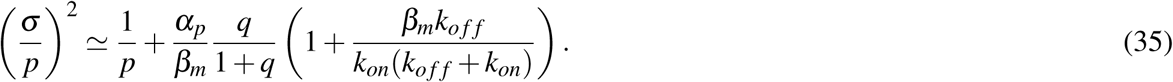

Using Eq. 29 which expresses transcription *β*_*m*_ as function of gene (in)activation parameters *k*_*on*_, *k*_*off*_, δ, we can eliminate *β*_*m*_ from the last term:

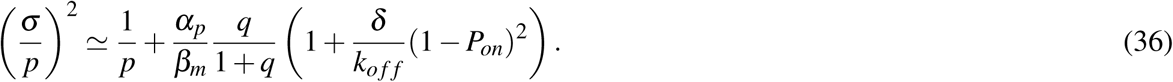

We also eliminate *δ* by introducing the transcriptional burst size *b*, which is the average number of mRNAs that are synthesized each time the gene is activated:

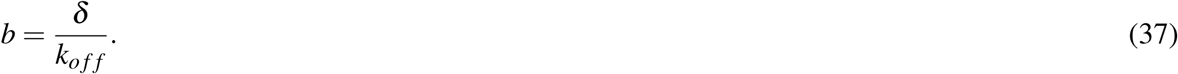

Plugging Eq. 37 for *b* into Eq. 36 for 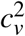, we obtain:

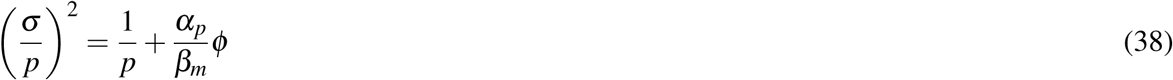

where

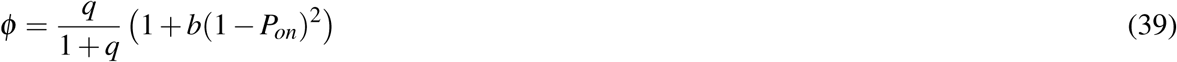

accounts for gene activation dynamics.

Except for highly expressed genes, the transcriptional burst size *b* is typically small (*b ≃* 1)^60^. For *q* = 3 (Table S1), varying *P*_*on*_ across its full range causes *ϕ* to vary only between 0.75 and 1.5 (Fig. S3E). Hence, in the worst case, neglecting gene activation dynamics by setting *ϕ* = 1 would result in a 1.5-fold error on 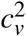. We conclude that neglecting gene activation dynamics by setting *ϕ* = 1 in Eq. 38 for 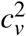 results in a reasonable approximation of the coefficient of variation for most genes, except for highly-expressed genes.

For highly-expressed genes, experiments in *S. cerevisiae*^50^ and *H. sapiens*^13^ found a noise floor. This noise floor might occur when the transcription rate exceeds the maximal rate of gene activation^13^. In this scenario, high transcription rate can only be achieved by increasing the burst size *b*^13,60^. Since *b* is not a constant in the large *β*_*m*_ regime, we rewrite *b* as in terms of the transcription rate *β*_*m*_. To do so, we combine Eq. 29 which expresses *β*_*m*_ in terms of the kinetic rates of gene activation (*k*_*on*_, *k*_*off*_, *δ*) and Eq. 37 which defines the burst size *b* in terms of *δ* and *k*_*off*_ to find

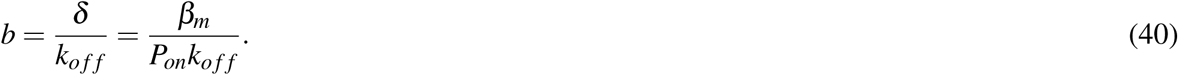

Plugging in this expression for *b* in Eq. 38 eliminates the burst size:

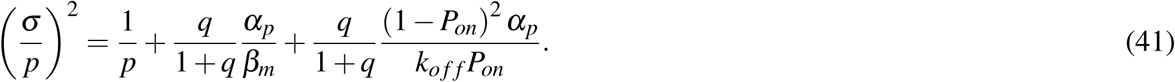

Neglecting the 1*/p* term which is in the order of 10^−3^ or smaller, we can get an expression for the noise floor *c*_*v*0_ by taking the limit of large *β*_*m*_. In this limit, the second term in Eq. 41 vanishes and 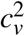 approaches

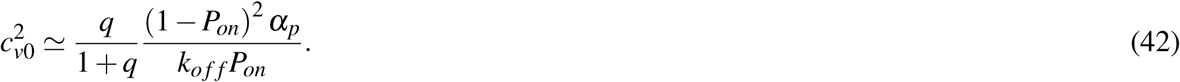

Plugging in gene activation parameter values typical for *H. sapiens*^13^ (P_on_ = 0.18, *k*_*off*_ = 2*h*^−1^) and setting *q* = *α*_*m*_/*α*_*p*_ = 3, *α*_*p*_ = 0.05*h*^−1^ (see Table 1) puts the noise floor *c*_*v*0_ at 0.27, a value comparable to experimental observations^13^ (*c*_*v*0_ ≃ 0.3). Hence, the noise floor on protein abundance could occur when transcription rates saturate gene activation kinetics. In this case, substituting Eq. 42 for 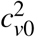 into Eq. 41 leads to:

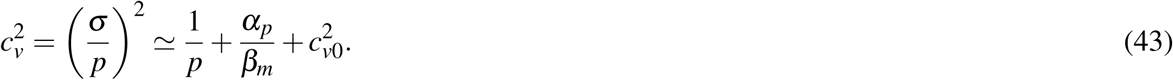

This final expression is identical to the one we previously derived for *E. coli* (Eq. 32). It would also hold if the noise floor was caused by a mechanism different from the transcriptional saturation of gene activation dynamics, such as extrinsic noise. Note that we neglected the *q/*(1 + *q*) term of Eq. 41. This is because the resulting approximation error on 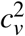 would be less than 50% (Fig. S3E). Because *β*_*m*_ scales inversely with 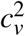 (Eq. 43), neglecting gene activation kinetics implies at most a 50% error on *β*_*m*_, or equivalently a 0.2 error on log^10^ *β*_*m*_. This is small compared the dynamic range of transcription rates which vary over 2–3 order of magnitudes.

### Power-law scaling of mRNA Fano factor with mRNA abundance cannot explain the noise floor

In *E. coli*, So et al. ^72^ observed that the Fano factor of mRNA scales as a power-law of mRNA abundance *m*,

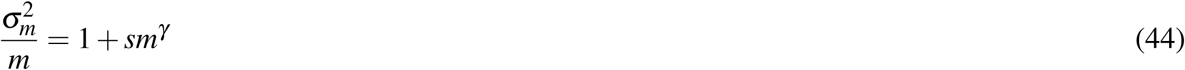

with *γ ≃* 0.64 and *s ≃* 1.5. This scaling holds for mRNAs whose abundance range from 0.3 to 40 to per cell^72^. A similar scaling was observed in a human fibroblast cell line^11^.

Here we ask whether this scaling can predict the noise floor, which would remove the need for the phenomenological factor 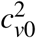 in the expression for the protein noise *c*_*v*_ derived in the previous section (Eq. 32).

In the case of the stochastic model of gene expression of the previous section (Fig. S3C), Paulsson ^54^ showed that the Fano factor for mRNA abundance *m* can we written as

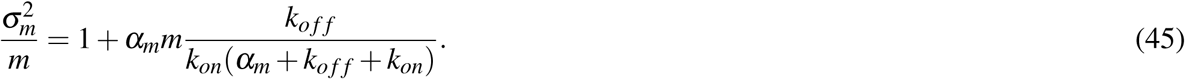

By equating the observed Fano factor power-law (Eq. 44) and the theoretical expression for the Fano factor (Eq. 45), we find how the gene activation noise term varies with the transcription rate *β*_*m*_:

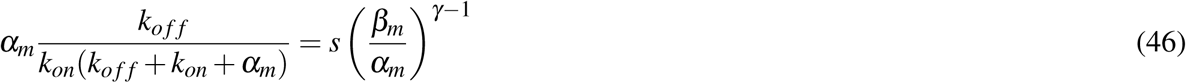

where we have used *m* = *β*_*m*_/α_*m*_. Plugging this expression into the expression for 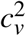 derived by Paulsson ^54^ (Eq. 28), we can find how protein noise 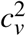 varies with transcription rate *β*_*m*_

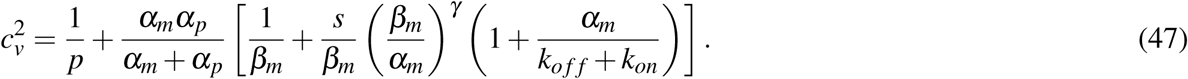

For *γ <* 1 and in the typical case in which gene (in-)activation is fast compared to mRNA decay *k*_*off*_, *k*_*on*_ » α_*m*_, 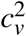 goes to 0 for large *p* and *β*_*m*_. Thus, there is no noise floor in this regime.

In the opposite regime where mRNA decay is fast compared to gene (in-)activation *α*_*m*_ *» k*_*off*_ + *k*_*on*_, we could have *k*_*on*_ ∼ *β*_*m*_ or 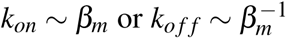. In the first case, 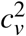 goes to 0 for large *p* and *β*_*m*_, so there is no noise floor. We already studied the second case in the previous section on *E. coli* noise and found that the resulting gene activation noise is too small compared to the observed noise floor. Finally, plotting 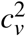 as a function of *β*_*m*_ for typical values of *α*_*m*_, *α*_*p*_, *s, k*_*on*_, *k*_*off*_ and *γ*, we confirmed that the Fano factor power law could not explain the noise floor observed in single cell experiments (not shown).

In conclusion, incorporating the Fano factor power law (Eq. 44) into the model, we find that it results in no noise floor for *γ* ≃ 0.5 or in a noise floor that is too low compared to experimental observations. Since the noise floor is a well-founded experimental observation^13,50,71,76^, the power law by itself cannot explain the full noise behavior. Additional biology, such as extrinsic noise or increased transcriptional bursting at large transcription rates, is needed to understand the noise floor.

### Genes dominated by comparable requirements of precision are predicted to share the same translation / transcription ratio

We consider a protein of abundance *p*. The protein contributes a quantity *f* (*p*) to the organism’s fitness. The cost of transcription Δ *f*_*m*_ is linear in the transcription rate *β*_*m*_, the (pre-)mRNA length *l*_*m*_ and the fitness cost of transcription per nucleotide *c*_*m*_,

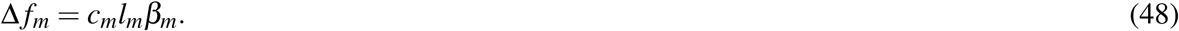

The overall fitness is thus *F* = *f* (*p*) *-* Δ *f*_*m*_. The fitness function *f* (*p*) reaches its maximum *f*_*max*_ at *p* = *p\**(Fig. 4A). Expanding *f* (*p*) around the optimum *p* = *p\** to second order, the overall fitness becomes

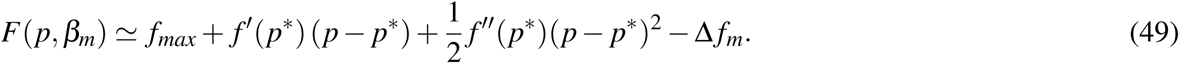

*f ^′^* (*p\**) = 0 since *f* (*p*) reaches its optimum *f*_*max*_ at *p* = *p\**. *f ^″^* (*p\**) is the curvature of the fitness function at its maximum. It is a negative number which characterizes how narrow the fitness function is.

Because protein abundance fluctuates around *p\**, the cell doesn’t experience the maximum fitness *f*_*max*_ but rather a lower average fitness *< f >*. Averaging *F* over fluctuations in protein abundance, we obtain

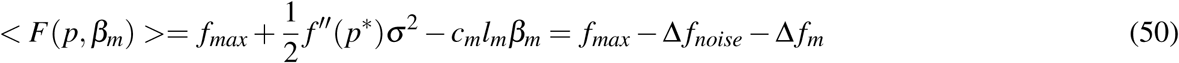

where *σ* ^2^ is the variance of protein abundance fluctuations (Fig. 4A). The curvature times this variance, 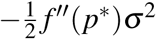 is the noise load Δ *f*_*noise*_, the fitness lost due to the stochastic fluctuations in protein abundance^32,81^ (Fig. 4A). It is a positive quantity because the curvature *f ^″^* (*p*) is negative at the maximum *p* = *p\**.

We seek the transcription rate *β*_*m*_ that maximizes fitness. For this purpose we note that *β*_*m*_ affects the noise level *σ* ^2^ in a well-characterized way. Theory and experiments of gene expression noise^5,50,54,76^ indicate that variance of protein noise is given by

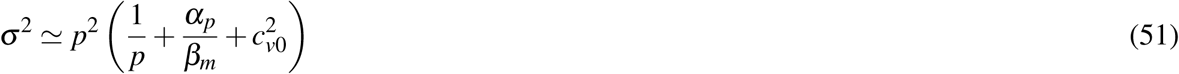

where *α*_*p*_ is the protein decay rate and *c*_*v*0_ is the noise floor due to extrinsic noise^76^ or the larger transcriptional burst size of high abundance proteins^13^ (see previous section). We can now solve for the *β*_*m*_ that maximizes fitness, by finding 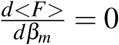. The optimal translation / transcription ratio rate *β*_*p*_/*β*_*m*_ satisfies

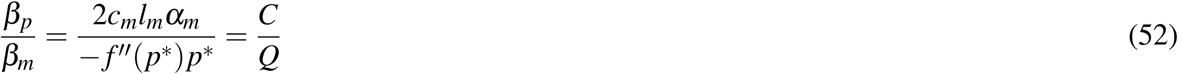

where we have used 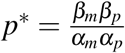. Note that we introduced two new variables *C* and *Q*.

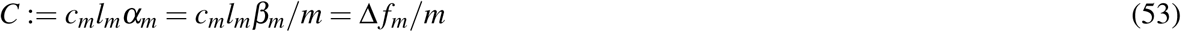

represents the fitness cost of transcription per mRNA molecule for a gene with pre-mRNA length *l*_*m*_.

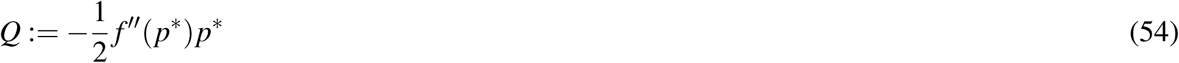

is the gene’s sensitivity to noise. Genes with narrower fitness functions (*f ^″^* (*p\**) *≪* 0) are more noise sensitive. A gene’s sensitivity to noise also depends on *p\** because stochastic fluctuations *σ* ^2^ scale with *p\** (Eq. 51).

From the expression for the optimal *β*_*p*_/*β*_*m*_ (Eq. 52), we see that genes with narrow fitness function have lower translation / transcription ratios (Fig. 4B). Higher transcription cost per mRNA molecule — due to longer mRNAs *l*_*m*_ or rapid mRNA turn-over *α*_*m*_ or a scarcity of nucleotides leading to increased cost *c*_*m*_ — shifts the balance towards higher ratios. Note that the translation / transcription ratio does not depend on translation cost, although this cost is typically larger than transcriptional cost^31,44,82^. This is because we assumed that, for a given protein abundance, translation cost are the same if the proteins are synthesized from few or many mRNAs.

### The lowest translation / transcription ratio can be predicted from measurable, fundamental parameters of each organism

In this section, we ask what is the predicted offset of the line that forms the boundary of the depleted region, namely the constant *k* such that *β*_*p*_/*β*_*m*_ > *k* for all genes. We provide estimates based on known fundamental parameters of cell biology suggested by the theory.

To estimate *k*, we note that the precision - economy theory predicts that low *β*_*p*_/*β*_*m*_ occur for genes with narrow fitness functions (Eq. 52, Fig. 4B). But the noise floor *c*_*v*0_ (Fig. 4C) sets a limit on how narrow fitness functions can be: for fitness functions that are too narrow given the noise floor, average fitness is negative (Fig. 4D). Such fitness functions cannot be selected for in evolution. We can therefore estimate the maximal transcription rate by determining the largest, selectable fitness function curvature, and then compute the optimal transcription rate for that function. The curvature should be small enough that the smallest, unavoidable protein fluctuations (set by the organism’s noise floor *c*_*v*0_) do not dominate the fitness benefit of expressing the protein. In other words, the fitness benefit of expressing the protein *f*_*max*_ should be larger than the noise load Δ *f*_*noise*_:

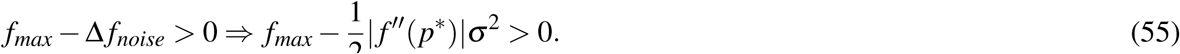

Here we neglected mRNA cost because for proteins with narrow fitness functions, it is small compared to the noise load Δ *f*_*noise*_.

If fitness is mainly set by the growth rate *µ*, the fitness contribution *f*_*max*_ of a gene cannot be larger than the growth rate: *f*_*max*_ *< µ*. In addition, fluctuations cannot be smaller than the noise floor, *σ/p\** > c_*v*0_ ^13,50,71,76^. From these two considerations, we can compute an upper bound on the noise sensitivity *Q*,

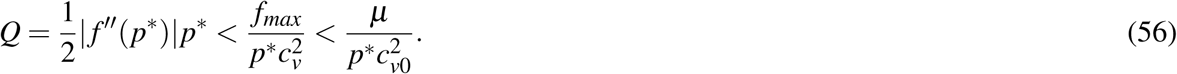

By substituting this upper bound in Eq. 52 for the optimal *β*_*p*_/*β*_*m*_, we find an upper bound on 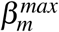 on transcription rates

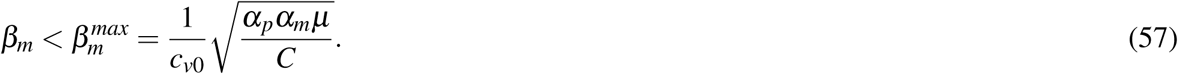

We now consider the (hypothetical) protein expressed at maximal transcription 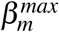 and maximal translation 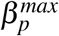. This protein has highest protein abundance *p\**. It also has narrowest fitness function (narrower fitness are not selectable due to the noise floor), and thus highest noise sensitivity *Q*_*max*_. We can plug Eq. 57 for *β*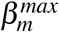 into Eq. 52 for the optimal *β*_*p*_/*β*_*m*_ to find *Q*_*max*_:

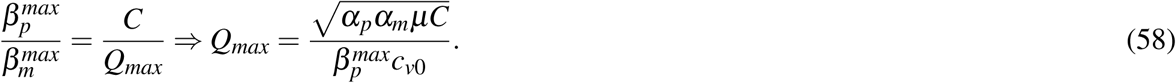

In this expression, we can see that a higher noise floor implies that genes need to be less sensitive to noise. We now use *Q*_*max*_ in the Equation 52 for the optimal *β*_*p*_/*β*_*m*_ to find *k*,

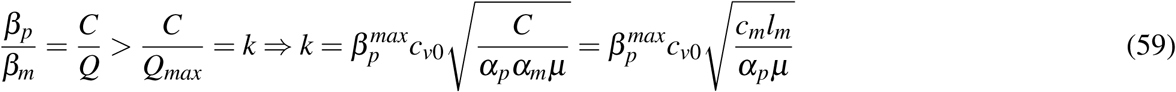

where we have used *C* = *c*_*m*_*l*_*m*_α_*α*_ (Eq. 53). The fitness cost per transcribed nucleotide *c*_*m*_ can be estimated from the average contribution of each nucleotide of each mRNA to the organism’s fitness, *c*_*m*_ = *µ/* Σ *β*_*m*_*l*_*m*_ (Eq. 13). Neglecting differences in mRNA length between genes, we finally find:

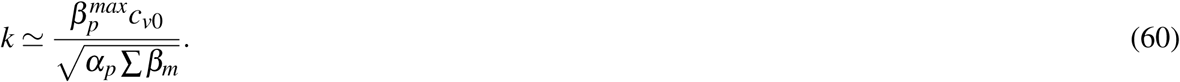

In this derivation, we have assumed that the gene’s contribution to fitness *f*_*max*_ is smaller than the growth rate *µ, f_max_ < µ* (Eq. 56). For essential genes, *f*_*max*_ ≃ *µ*. For non-essential genes, we can use a tighter upper-bound on *f*_*max*_:

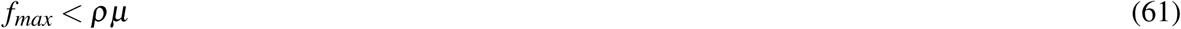

with 0 *< ρ <* 1. For example, if deleting a gene decreases fitness by 1% or less, we have *ρ* = 0.01. By repeating the derivation, we find

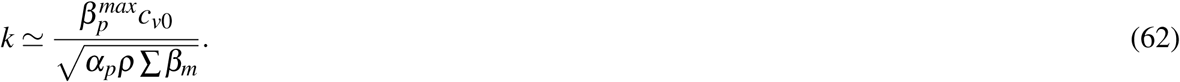

Thus, for non-essential genes (*ρ «* 1), the predicted boundary of the depleted region has higher intercept.

### Estimating the curvature of fitness functions from the measurements of Keren et al

Keren et al. ^34^ measured the fitness of *S. cerevisiae* cells as a function of log-expression for growth in glucose. Specifically, these measurements map log_10_ gene expression *x* to fitness *f* (*x*), with *x* = log_10_ *p*. These measurements do not allow to estimate the curvature of the fitness function *f ^″^* (*p\**) directly. Nevertheless, they allow to estimate a closely related quantity, namely:

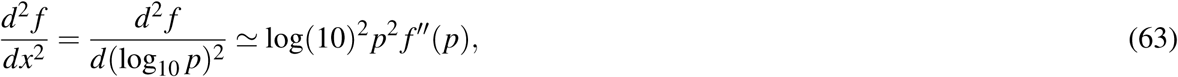

where we neglected *f ^′^* (*p*) since we expand around the fitness optimum. In this section, we compare the measured *p*^2^ *f ^″^* (*p*) to the predictions from the theory.

We focused on genes present in both the study of Keren et al. ^34^ and the ribosomal profiling data of Weinberg et al. ^84^. To have enough context to estimate the local curvature, we considered only genes that could be both under-expressed and over-expressed compared to their wild-type expression by at least half an order of magnitude. Following Keren et al. ^34^, we also discarded low-quality genes, such as genes whose fitness value at wild-type expression was significantly lower than the fitness of the wild-type. This left 34 genes for analysis. We further excluded 9 genes with TATA promoters because these genes tend to have large transcriptional burst size^88^: our model assumes a small transcriptional burst size and hence cannot accurately model the noise of genes with TATA promoters. Finally, we did not consider 4 genes whose curvature was too low to be estimated accurately given the accuracy of fitness measurements (*| f ^″^* (*p\**)*p\**^2^ *<* 0.006). This leaves 21 genes for the analysis.

To estimate the local curvature at wild-type log_10_ expression *x*_*wt*_, we focus on fitness measurements located within one order of magnitude of *x*_*wt*_ : *x*_*wt*_ - 1 *< x < x_wt_* + 1.

To these measurements, we then fit the parameters *a, c, d* of the polynomial:

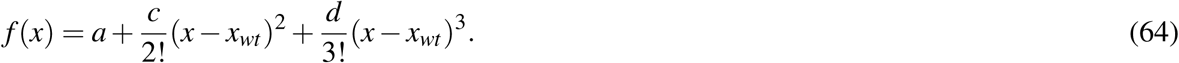

There is no first order term because fitness peaks at wild-type expression. The third order term allows for asymmetry in the fitness function. *c* is the curvature of *f* (*x*) at *x*_*wt*_. Using Eq. 63 for the curvature of fitness as a function of log_10_ *p*, we find a relationship between the curvature and the *c* parameter of the polynomial:

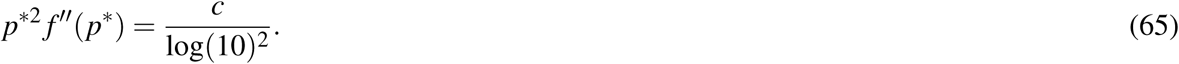

We estimate the standard error on *c* using Fisher’s information matrix, assuming a 10% error on fitness measurements^34^.

By rewriting Eq. 52 for the optimal *β*_*p*_/*β*_*m*_ in terms of *f ^″^* (*p\**)*p\**^2^, we can predict the local curvature of fitness functions from the precision - economy theory:

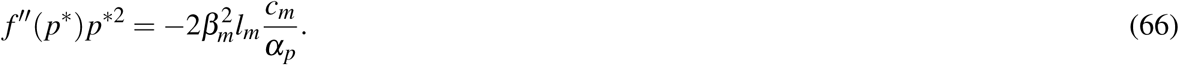

We estimate the error on these predictions by considering that the sampling error on log_10_ *β*_*m*_ is about 0.1 (see section on data processing), and a 2-fold error on *c*_*m*_ (mainly due to uncertainty in the number of mRNAs per cell). This leads to a standard error of 0.36 on the log_10_ predictions.

We find that predictions and measurements of | *f* ^*″*^ (*p\**)|*p\**^2^ span the same range (10^−3^ – 10^−1^). The correlation coefficient between predictions and measurements is positive (*r* = 0.39).

To determine the significance of the agreement between predictions and measurements, we compute the root-mean-square deviation (RMSD) between predictions and measurements. Upon shuffling the measurements 10^6^ times, only in *p* = 4.1% of shuffles is the RMSD smaller than the RMSD computed on the non-shuffled measurements. Hence, the agreement between predictions and measurements is unlikely explained by chance. Consistent with this result, a *χ*^2^ test concludes that predictions and measurements do not differ significantly given the measurement error (*p* = 0.51).

### The region in which transcription noise is comparable to the noise floor does not correspond to the depleted region

The depleted region cannot be explained by determining the *β*_*m*_ and *β*_*p*_ for which increasing transcription provides little extra precision relative to the the noise floor.

To see why, consider Eq. 32 which relates transcription *β*_*m*_ to the noise *σ/p*:

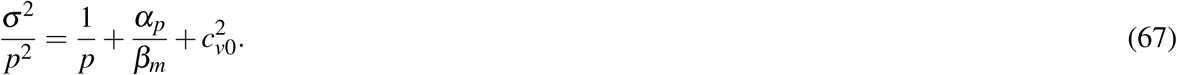

Transcription noise becomes comparable to the noise floor when 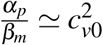. Thus, if the depleted region corresponds to combinations of *β*_*m*_ and *β*_*p*_ for which transcription provides little extra precision compared to the noise floor, the boundary of the depleted region should be

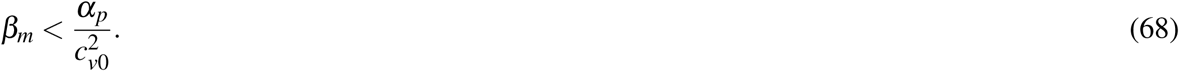

Therefore, the predicted boundary is a vertical line in the Crick space. This is a bad model for the data. For example, in *S. cervisiae*, the predicted boundary would be log_10_ *β*_*m*_ = 1.6 which excludes about half of the genome (Fig. 2A).

### Identifying groups of genes with significantly high or low translation / transcription ratio

Do genes with high or low translation / transcription ratio have different biological functions? To find out, one could iterate through groups of genes of similar function — as defined by Gene Ontology (GO) annotations^2^ — and test statistically whether genes in these groups show significantly high or low *β*_*p*_/*β*_*m*_. However, this approach neglects a key property of the central dogma rates: high abundance proteins cannot have high translation / transcription ratios due to the maximal translation rate 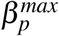. Since translation rates *β*_*p*_ cannot exceed 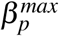, achieving high protein abundance requires recruiting transcription, which decreases *β*_*p*_/*β*_*m*_.

As a result, stratifying genes by equal *β*_*p*_/*β*_*m*_ arbitrarily groups low abundance proteins with low *β*_*p*_/*β*_*m*_ together with high abundance proteins that would have high *β*_*p*_/*β*_*m*_ but cannot because translation rates cannot exceed *β*_*p*_.

To address this issue, we stratify genes by their position relative to two boundaries: the line of lowest possible translation / transcription ratios (*β*_*p*_ = *k*β*_m_*), and the line of highest possible translation rates 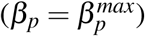. We summarize the position of genes in between these two boundaries by an angle *θ* (Fig. S5C). Genes that sit on the line of lowest possible translation / translation ratios have *θ* = 0, whereas *θ* = *π/*4 corresponds to genes with maximal translation / transcription ratios.

*θ* can be computed from a gene’s *β*_*m*_ and *β*_*p*_, and from the maximal transcription and translation rates 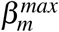 and *β* 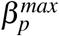 by trigonometry (Fig. S5C):

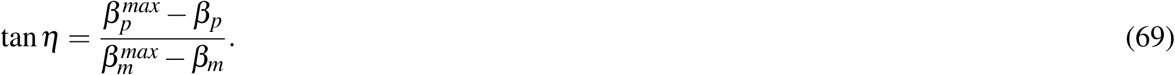

The line of lowest possible translation / transcription ratio has slope 1 (*β*_*p*_ = *k*β*_m_*), *η* + *θ* = *π/*4. Therefore,

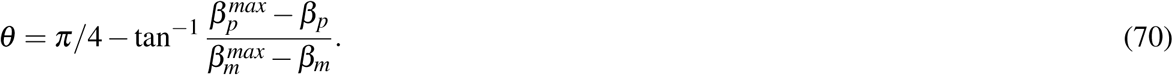

For each group of genes from the Gene Ontology, we use the Mann-Whitney test to compute the *p*-value that the *θ* s in that group are significantly larger or smaller compared to the distribution of *θ* s in genes overall. We estimate the false discovery rates (*q*-values) using the fdrtool R package^74^.

For all four model organisms, Supplementary Item 1 lists all GO categories with significantly high or low *θ* with false discovery rate *q <* 0.01, together with (uncorrected) *p*-values and *θ* normalized to *π/*4 so that *θ* ranges between 0 (for GO categories with lowest *β*_*p*_/*β*_*m*_) and 1 (for GO categories with highest *β*_*p*_/*β*_*m*_). Table S2 shows a summary of these results. Fig. S5D–G shows the position of genes belonging to selected significant GO categories in the Crick space.

### Differences in the genes groups with high and low *β*_*p*_/*β*_*m*_ in HeLa compared to 3T3 cells can be explained by global differences in gene expression profiles

Different groups of genes take unusually high or low *β*_*p*_/*β*_*m*_ in HeLa cells compared to 3T3 cells (Table S2). Here we examine whether these differences can be explained by global differences in the gene expression.

We obtained log^2^ RPKMs for HeLa and 3T3 cells the RNAseq experiments of Eichhorn et al. ^16^. Because 3T3 cells are of murine origin, comparing the two cell lines requires mapping mouse gene names to human names. To so so, we used ENSEMBL’s orthology database, using only one-to-one orthology relations of highest confidence (confidence = 1). After mapping genes names from mouse to human, we discarded 31 non-unique genes.

To provide more context for the gene expression analysis, we merged the HeLa and 3T3 gene expression profiles with the Cancer Cell Line Encyclopedia (CCLE) gene expression data^6^ based on human gene names. We excluded haematopoietic and lymphoid cell lines which form a distinct group of liquid tumors compared to the majority of solid tumor cell lines in this dataset. We quantile normalized the gene expression data of all the cell lines together.

To visualize the position of HeLa and 3T3 cell lines relative to other cancer cell lines, we performed principal component analysis on the log gene expression matrix of 859 cell lines x 6300 genes. The position of the different cell lines (gray dots) in the gene expression space spanned by first 3 PCs is shown on Fig. S5H.

Finally, we determined gene sets that were differently regulated in HeLa compared to 3T3 by gene set enrichment analysis^75^ (Fig. S5I).

---

**Figure S1.**
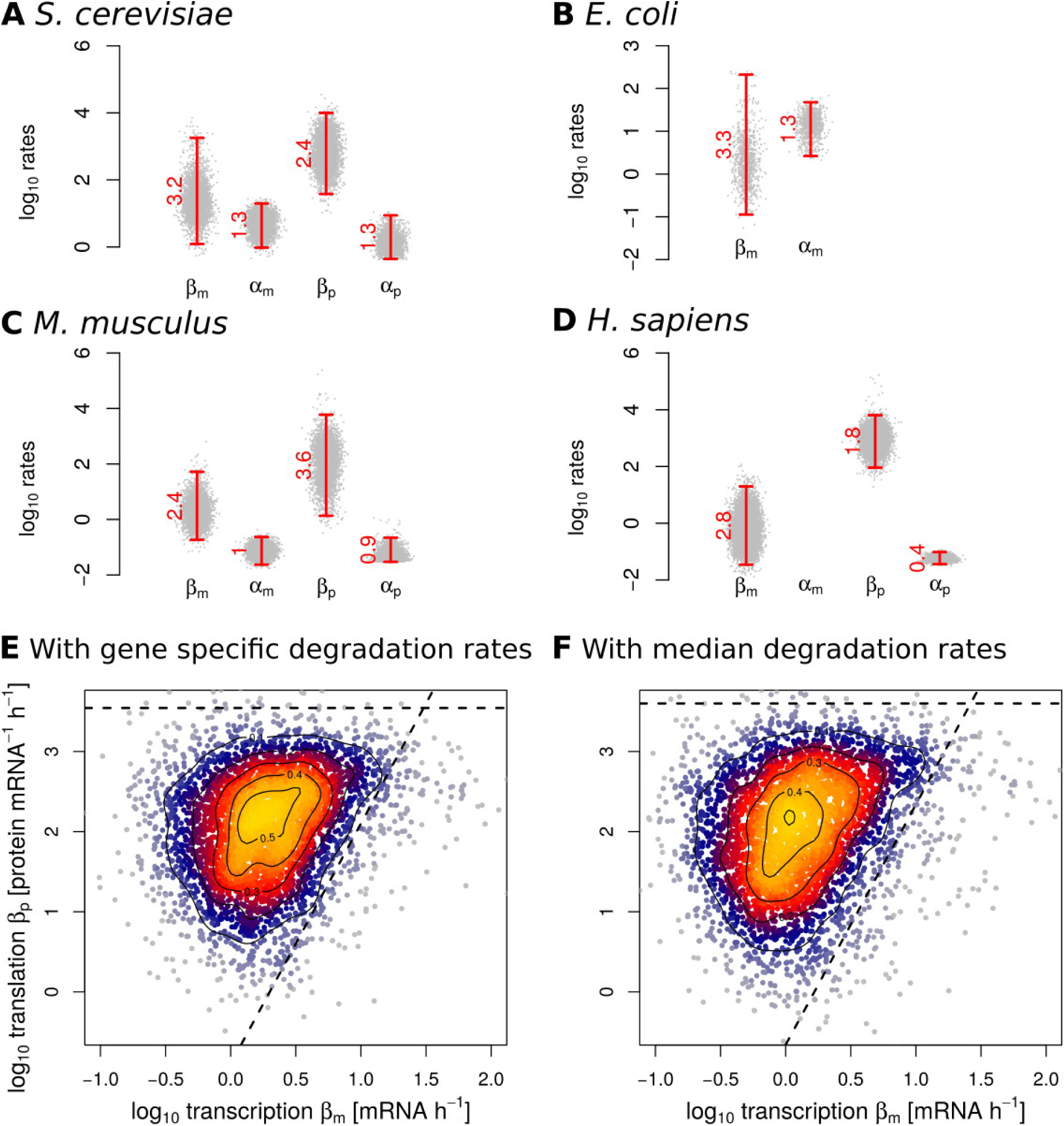
Related to Figure 1. **A – D.** Distribution of transcription, translation and mRNA and protein decay rates in four model organisms. **A.** *S. cerevisiae*: transcription and translation rates from Weinberg et al. ^84^, mRNA decay rates from Eser et al. ^18^, protein decay rates from Belle et al. ^7^. **B.** *E. coli*: transcription rates from Li et al. ^41^, mRNA decay rates from Chen et al. ^11^. **C.** *M. musculus*: all rates from Schwanhäusser et al. ^66^. **D.** *H. sapiens*: transcription and translation rates from Eichhorn et al. ^16^, protein decay rates from Cambridge et al. ^9^. Rates were estimated as described in the Methods. Bars span the range from the 0.5% quantile to the 99.5% quantile of rates (i.e. 99% of genes). **E – F.** Taking into account the specific mRNA and protein decay rates of each gene has a negligible effect on the distribution of genes in the Crick space. **E.** Transcription and translation rates as estimated from measurements of mRNA and protein abundances and decay by Schwanhäusser et al. ^66^. **F.** Same as E., except that transcription and translation rates where estimated from mRNA and protein abundance by setting mRNA and protein decay rates to their median. Gene positions in panels E and F differ by 0.3 (root mean square deviation in log^10^ rates), which is small (10%) compared to the dynamic range of transcription and translation rates which vary over three orders of magnitude.

**Figure S2.**
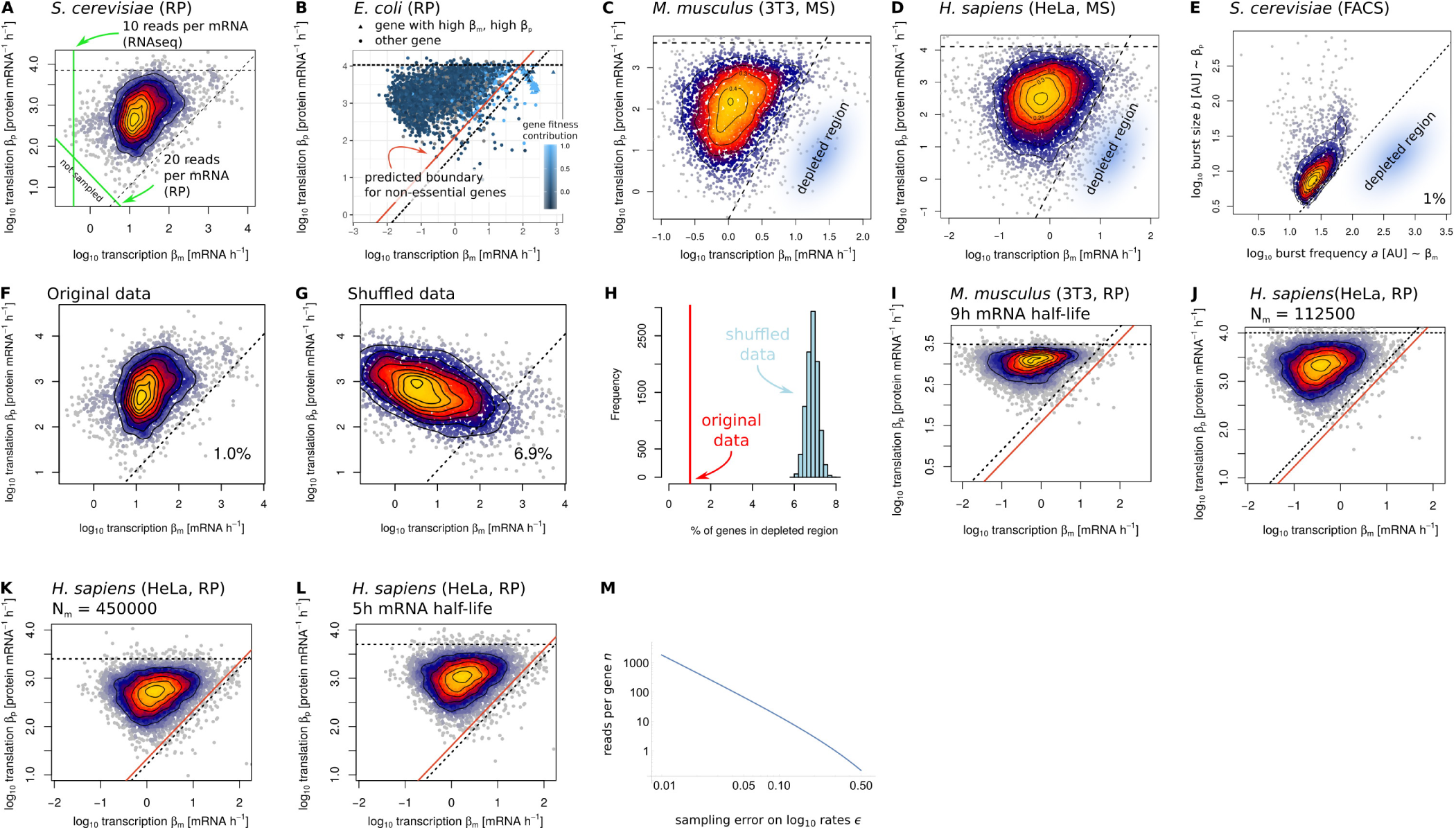
Related to Figure 2. **A.** The ‘not sampled’ region is explained by sequencing depth and our focus on genes with at least 10 mRNAseq reads and 20 ribosome profiling (RP) reads per mRNA (two green lines). **B.** In *E. coli*, a group of 62 genes (triangles) contribute strongly to fitness. Excluding these genes, the boundary of the depleted region (diagonal dotted black line) has intercept log^10^ *k* = 1.7 *±* 0.1, which is higher than log^10^ *k* = 1.1 *±* 0.1 found using all genes. A higher intercept is expected for non-essential genes (Methods). This is illustrated by the red line, which represents the boundary predicted for genes that contribute 1% of the organism’s fitness. Fitness data: Baba et al. ^3^. **C – E.** The depleted region is found in the mass spectrometry (MS) and mRNAseq data of Schwanh ä usser et al. ^66^ (panel C) and Nagaraj et al. ^49^ (panel D), as well in the flow cytometry data of Newman et al. ^50^ (panel E). By estimating transcription and translation rates from the mean protein abundance and coefficient of variation^22,23^, we find a depleted region whose boundary has slope 1 (panel E), as in ribosome profiling datasets. The boundary of the depleted region in the two proteomics datasets has slope larger than one 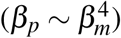 The different slope observed with mass-spectrometry compared to ribosome profiling and flow cytometry datasets could be explained by technical limitations in these pioneering mass-spectrometry datasets and differences in the error structure of mRNAseq and mass-spectrometry datasets^42^. **F – H.** Genes with high *β*_*m*_ and low *β*_*p*_ are statistically depleted. **F.** The boundary of the depleted region is a line of slope 1 such that 99% of genes are above the line. *S. cerevisiae* data from Weinberg et al. ^84^. **G.** Shuffling *β*_*m*_ and *β*_*p*_ while keeping the marginal distributions of *β*_*p*_ and protein abundance constant increases the fraction of genes located in the depleted region. **H.** All 10^4^ re-shuffled sets of *β*_*m*_ and *β*_*p*_ show a increased number of genes in the depleted region. Thus, the depletion of this region is statistically significant (*p <* 10^−4^). Repeating this procedure in *E. coli, H. sapiens* and *M. musculus* leads to the same conclusion (*p <* 10^−4^, not shown). **I – L.** The depleted region and the predicted boundary of this region are robust to uncertainties in mRNA half lives and in total number of mRNAs per cell *N*_*m*_. **M.** The error on log^10^ mRNA abundance *ε* can be controlled by discarding genes with low number of reads *n*.

**Figure S3.**
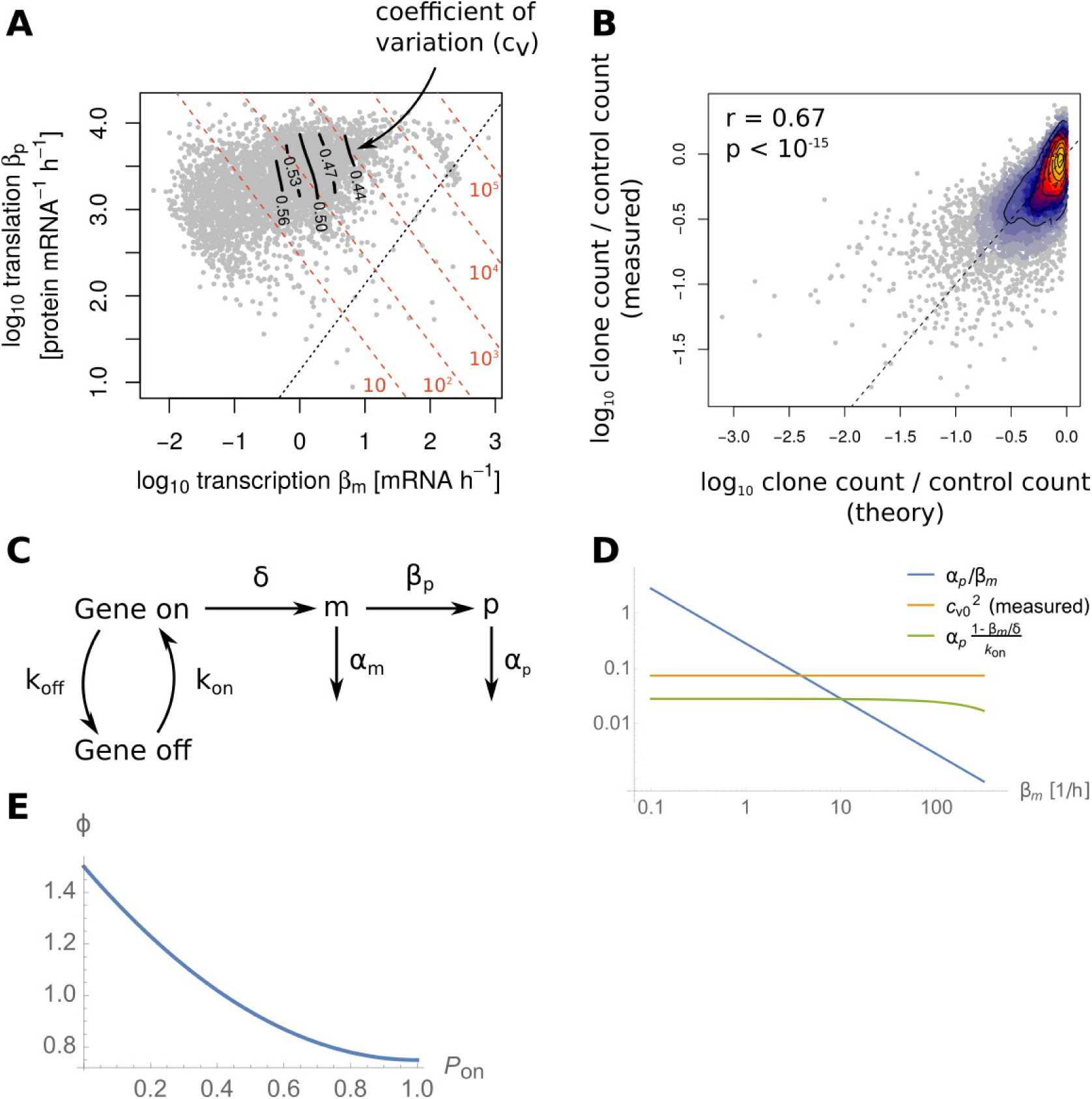
Related to Figure 3. **A.** In *E. coli*, coefficients of variation scale with transcription rates. Transcription and translation rates inferred from Li et al. ^41^. CVs from Taniguchi et al. ^76^. **B.** Clone abundance in growth competition experiments can be predicted from theoretical estimates of fitness cost of mRNA *c*_*m*_ and fitness cost of protein *c*_*p*_. We assumed a growth time *t* = 24h in this figure. The correlation between predictions and measurements is independent of the growth time *t* (Methods). The dotted line represents a perfect fit between measurements and experiments (*y* = *x*). **C.** The 3-stage telegraph process can be used to models how intrinsic noise in gene expression is affected by different reaction rates. **D.** In the 3-stage model of gene expression of panel C, intrinsic protein noise is mainly due to small mRNA copy number noise (blue line) when *β*_*m*_ is small. For large *β*_*m*_, intrinsic protein noise is dominated by gene activation noise (green line), which is almost constant for *β*_*m*_ in physiological range (x-axis). Constants are from the experiments of So et al. ^72^ and Taniguchi et al. ^76^ : *α*_*p*_ = 0.28/h, 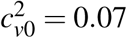, *δ* = 800/h, *k*_*on*_ = 10/h. With these constants, the noise floor observed experimentally (orange line) is larger than the gene activation noise. **E.** By neglecting gene activation dynamics, one can estimate the coefficient of variation from the central dogma rates with an error *ϕ* less than 1.5-fold. Note however that this result does not hold for highly expressed genes (see Methods).

**Figure S4.**
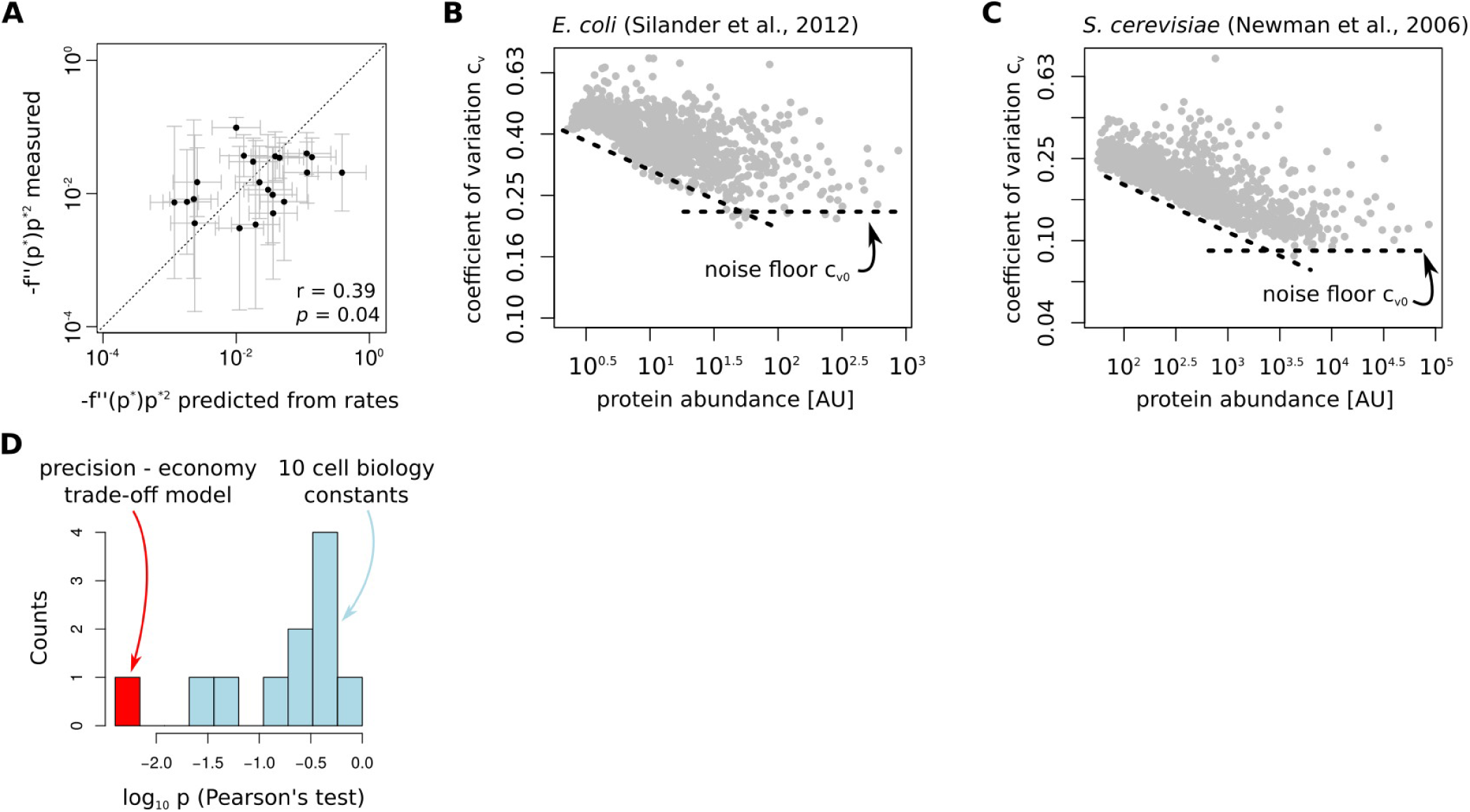
Related to Figure 4. **A.** Measured curvatures of fitness functions are within measurement error of curvatures predicted by theory from transcription and translation rates. Curvatures were estimated from the experiments of Keren et al. ^34^ who measured fitness as a function of log protein abundance. The quantity *f ^″^*(*p\**)*p\**^2^ can be directly estimated from this data (Methods). We used the precision -economy theory to predict *f ^″^*(*p\**)*p\**^2^ from the central dogma rates (Methods). Error bars represent standard errors. **B.** A noise floor cv0 = 0.22 ±0.01 is found in the measurements of the protein noise conferred by *E. coli* promoters of Silander et al. ^76^. This is comparable to the noise floor *c*_*v*0_ = 0.27 *±* 0.01 found in the measurements of Taniguchi et al. ^76^ (Fig. 4C). **C.** A noise floor *c*_*v*0_ is also found in *S. cerevisiae*. Coefficients of variation and protein abundance from Newman et al. ^50^. **D.** Individual constants cannot predict the position *k* of the boundary of the depleted region. We correlated the position of the boundary of the depleted region with the predictions from the theory as well as with the 8 cell biology constants listed in Table 1 (number of mRNAs *N*_*m*_ and protein *N*_*p*_ per cell, cell volume *V*, growth rate *µ*, active protein decay rate *α*_*deg*_, effective protein decay *α*_*p*_, mRNA decay *α*_*m*_, noise floor *c*_*v*_0), in addition to the maximal translation rate 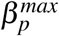 and total transcription output Σ *β*_*m*_ (Table 1). We tested for significant correlations between these constants and measured *k*. Only with the precision -economy theory do we find a significant correlation between measurements and theory (*p <* 0.01).

**Figure S5.**
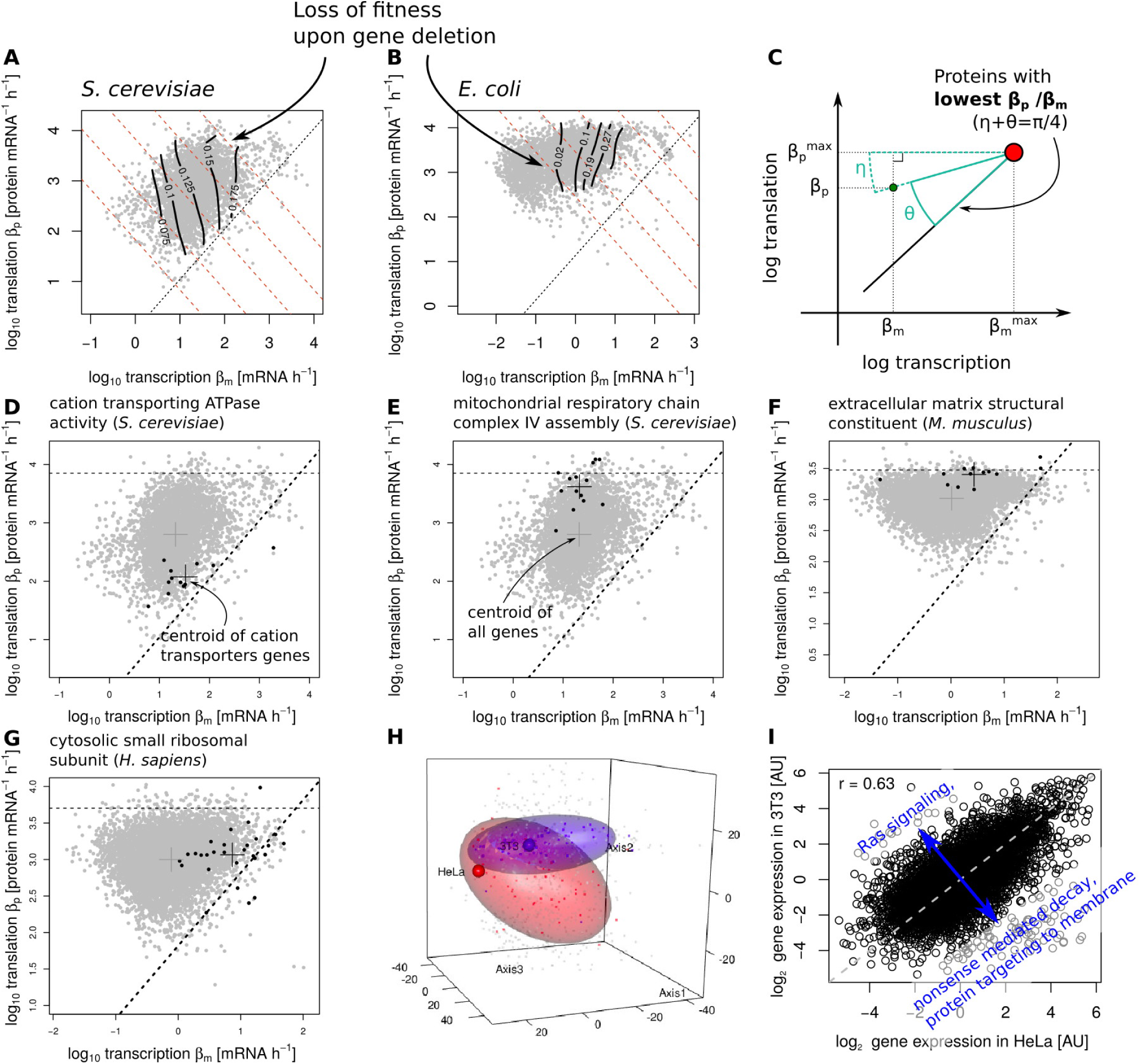
Related to Figure 5. **A – B.** Genes located close to the boundary of depleted region provide more fitness benefit than genes located far from it. Contours of fitness loss upon gene deletion in *S. cerevisiae* (panel A) and *E. coli* (panel B). We obtained fitness data from Steinmetz et al. ^73^ (*S. cerevisiae*) and Baba et al. ^3^ (*E. coli*). Fitness was Gaussian-smoothed using the same procedure as the CV data (Methods). **C.** To test for statistical associations between the function of genes and their position in the Crick space, we describe each gene by an angle *θ*. *θ* = 0 for genes with lowest possible translation / transcription, whereas genes that maximize the translation / transcription ratio have *θ* = *π/*4. *θ* can be computed from a gene’s *β*_*m*_ and *β*_*p*_, and from the maximal transcription and translation rates 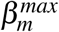 and 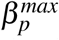. **D – G.** Genes with different functions (GO categories) are found close or far from the boundary of the depleted region. In each panel, black dots represent genes belonging to a specific group of genes. **H.** Principal component analysis of gene expression in 859 cancer cell lines (gray dots) shows that HeLa cells cluster with ovarian cell lines (red dots) whereas 3T3 cells cluster with skin cell lines (blue dots). Ellipses represent the covariance of skin and ovarian cell lines at the 75% confidence level. **I.** 3T3 cells over-express Ras signaling genes whereas HeLa cells over-express genes involved in nonsense mediated decay and protein targeting to membrane (Gene Set Enrichment Analysis, see Methods).

**Table S1.** Constants used to compute transcription and translation rates from mRNAseq, ribosome profiling and proteomics data. Listed are values for the number of mRNAs per cell *N*_*m*_, cell volume *V* [*µ*m^3^], number of proteins per cell *N*_*p*_, growth rate *µ* [h^−1^], protein degradation rate *α*_*deg*_ [h^−1^, protein decay rate *α*_*p*_ = *α*_*deg*_ + *µ* [h^−1^], mRNA decay rate *α*_*m*_ [h^−1^] and noise floor *c*_*v*0_. References for data sources are detailed in the Methods.

| Organism | $N_m$ | $V$ | $N_p$ | $\mu$ | $\alpha_{deg}$ | $\alpha_p$ | $\alpha_m$ | $c_{v0}$ |
| --- | --- | --- | --- | --- | --- | --- | --- | --- |
| *S. cerevisiae* | 60000 | 37 | $1.1 \times 10^8$ | 0.42 | 0.92 | 1.3 | 5.1 | 0.1 |
| *M. musculus* (3T3) | 180000 | 2000 | $6.0 \times 10^9$ | 0.03 | 0.01 | 0.04 | 0.14 | 0.3 |
| *H. sapiens* (HeLa) | 230000 | 2500 | $7.5 \times 10^9$ | 0.03 | 0.02 | 0.05 | 0.06 | 0.3 |
| *E. coli* | 1400 | 1 | $3 \times 10^6$ | 1.9 | 0 | 1.9 | 15 | 0.25 |

**Table S2.**
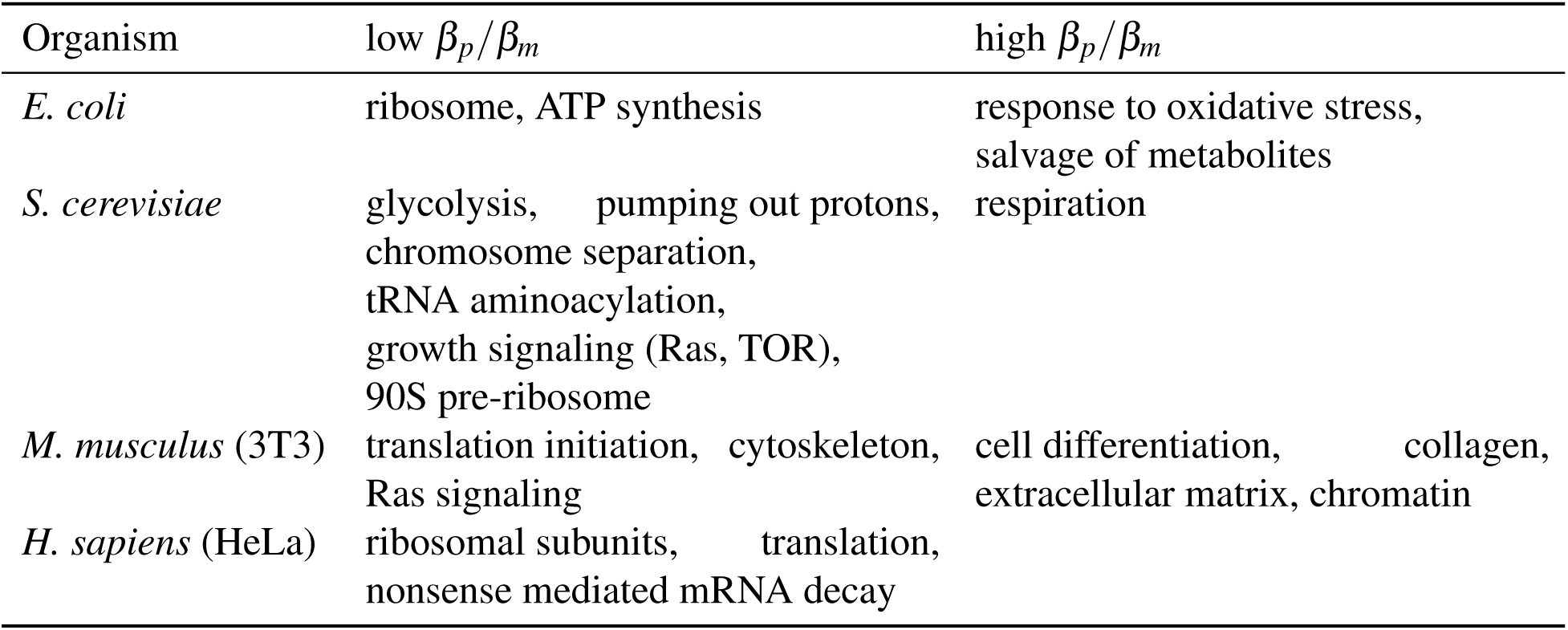
Genes that are key to growth tend have low *β*_*p*_/*β*_*m*_ whereas genes needed for stress response, survival and differentiation have high *β*_*p*_/*β*_*m*_. For each of the four organisms studied in the present study, the table lists GO categories that are significantly enriched (FDR *<* 0.01) at high and low *β*_*p*_/*β*_*m*_. The observation that respiration genes have high *β*_*p*_/*β*_*m*_ in *S. cerevisiae* could hint at a bet-hedging strategy, where a fraction of the population prepares for growth in a resource-limited environment. Comparing 3T3 and HeLa cells, we find that genes involved in translation initiation, ATP binding, RNA binding as well as ubiquitin-dependent protein catabolism appear both in both mouse and human cells (Supplementary Item 1). Other groups of genes are only found in one of the two cell lines. Such difference in GO terms is expected for cancer cell lines of different tissue origin: 3T3 cells cluster with skin cell lines whereas HeLa cells cluster with ovary cell lines (Fig. S5H, Methods). In addition, differences in genes expressed by 3T3 and HeLa cells can explain differences in genes with high and low *β*_*p*_/*β*_*m*_ (Methods). For example, 3T3 over-express Ras signaling genes (Fig. S5I). These genes have significantly low *β*_*p*_/*β*_*m*_ in 3T3 cells but not in HeLa (Supplementary Item 1). In contrast, HeLa cells over-express genes involved in nonsense mediated decay and SRP-mediated protein targeting to membrane (Fig. S5I) which have low *β*_*p*_/*β*_*m*_ in HeLa but not in 3T3 (Supplementary Item 1). Therefore, differences in GO terms with high and low *β*_*p*_/*β*_*m*_ in human and mouse can be explained by differences in the underlying biology of the cell lines.

